# miRNA family miR-29 inhibits PINK1-PRKN dependent mitophagy via ATG9A

**DOI:** 10.1101/2024.01.17.576122

**Authors:** Briana N Markham, Chloe Ramnarine, Songeun Kim, William E Grever, Alexandra I Soto-Beasley, Michael Heckman, Yingxue Ren, Andrew C Osborne, Aditya V Bhagwate, Yuanhang Liu, Chen Wang, Jungsu Kim, Zbigniew K Wszolek, Owen A Ross, Wolfdieter Springer, Fabienne C Fiesel

**Author notes:** Department of Medical and Molecular Genetics, Stark Neurosciences Research Institute, Indiana University School of Medicine, Indianapolis, IN 46202, USA. these authors contributed equally (Wolfdieter Springer, Fabienne C Fiesel).

## Abstract

Loss-of-function mutations in the genes encoding PINK1 and PRKN result in early-onset Parkinson disease (EOPD). Together the encoded enzymes direct a neuroprotective pathway that ensures the elimination of damaged mitochondria via autophagy.

We performed a genome-wide high content imaging miRNA screen for inhibitors of the PINK1-PRKN pathway and identified all three members of the miRNA family 29 (miR-29). Using RNAseq we identified target genes and found that siRNA against ATG9A phenocopied the effects of miR-29 and inhibited the initiation of PINK1-PRKN mitophagy. Furthermore, we discovered two rare, potentially deleterious, missense variants (p.R631W and p.S828L) in our EOPD cohort and tested them experimentally in cells. While expression of wild-type ATG9A was able to rescue the effects of miR-29a, the EOPD-associated variants behaved like loss-of-function mutations.

Together, our study validates miR-29 and its target gene ATG9A as novel regulators of mitophagy initiation. It further serves as proof-of-concept of finding novel, potentially disease-causing EOPD-linked variants specifically in mitophagy regulating genes. The nomination of genetic variants and biological pathways is important for the stratification and treatment of patients that suffer from devastating diseases, such as EOPD.

## INTRODUCTION

Parkinson’s disease (PD) is the second most common neurodegenerative disease. PD is clinically characterized by motor symptoms such as gait impairment, resting tremor, and muscular rigidity [1]. Pathologically, PD is characterized by dopaminergic neuron loss in the substantia nigra *pars compacta* and the consequential basal ganglia dysfunction [1]. Recessively inherited loss-of-function mutations in PTEN-induced kinase 1 (*PINK1*) and the E3 ubiquitin ligase *PRKN* are the most common genetic causes for early-onset PD, which occurs in about 5-10% of cases [1,2]. PINK1 and PRKN are at the center of a mitochondrial quality control pathway that eliminates damaged mitochondria through selective autophagy [3,4], suggesting that this mitophagy pathway is crucial for the survival of dopaminergic neurons. Originating from observations of mitochondrial toxins causing parkinsonism [5], a role for mitochondrial stress had long been implicated in the pathogenesis of PD.

Upon mitochondrial depolarization, PINK1 stabilizes on the outer mitochondrial membrane where it phosphorylates ubiquitin (Ub) and PRKN at Ser65 [6–11]. Both events are required for the full activation of the pathway. The translocation of cytosolic PRKN to mitochondria leads to the jointly mediated formation of mitochondrial pS65-Ub chains [12]. These serve as a specific signal for the engulfment of mitochondria in autophagosomes to facilitate their degradation by autophagy [13]. pS65-Ub has been shown to accumulate as cytoplasmic granules during aging, stress, and disease, making it a distinctive feature of mitochondrial stress and damage [14,15] further supporting a role of PINK1-PRKN-dependent mitophagy for human disease.

Since PINK1-PRKN-dependent mitophagy is a critical cell biological pathway for the survival of dopaminergic neurons and because mutations in PINK1 and PRKN only explain a small portion of EOPD cases, we hypothesized that loss of function of other mitophagy regulating genes might also be involved in the etiology of EOPD. Multiple genome-wide siRNA screens have already identified an extensive number of mitophagy genes although with little overlap between studies [16–19]. In order to complement these data sets and to filter for high confidence mitophagy regulators, we set out to perform a genome-wide miRNA screen. We have previously shown that the PINK1-PRKN pathway can be modulated by miRNAs [20]. The goal of our study was to discover novel, regulatory genes of the PINK1-PRKN pathway which could play a role for the etiology of EOPD, similar to PINK1 and PRKN mutations.

## RESULTS

### Genome-wide miRNA screening of PRKN modulators

For genome-wide screening of miRNAs, we used an established stable HeLa cell model with homogenous expression of GFP-PRKN [21]. We first tested different concentrations of the mitochondrial depolarizer carbonyl cyanide m-chlorophenyl hydrazone (CCCP) for the screening of inhibitory miRNAs and selected 3.5 µM CCCP, which was sufficient to fully activate GFP-PRKN translocation (**Fig. S1A**). We then reverse-transfected the cells with a library containing 1902 miRNAs and treated with CCCP for 2 h. Successful transfection was confirmed by a killer siRNA [22] that reduced the cell count by 85-90% compared to negative control-transfected or untransfected cells. Images from the screen were scored by a linear classifier algorithm that was developed using guided machine learning. As a training set for machine learning, we used cells that were transfected with non-targeting siRNA and treated with DMSO or with CCCP for 2 h. 415 different imaging parameters (**Table S1**) were used to find the best descriptors for the GFP-PRKN translocation phenotype. The final classifier that was employed for the image algorithm used a combination of twelve parameters to detect PRKN translocation (**Table S2, S3 and Fig. S1B**). Using this readout, we found that 86 out of the 1902 miRNAs (4.5%) reduced PRKN translocation percentage to under 50% (**Fig. 1A, B**). The new linear classifier was in excellent agreement with our previously used GFP-PRKN ratio algorithm that determines PRKN translocation by measuring the ratio of cytoplasmic over nuclear GFP-PRKN signal [21,23–25] (**Fig. 1C**) but was more robust in this screening setting. The Z’ score with the machine-learning guided classifier was 0.79 and the Z’ of individual plates was larger than 0.6.

**Figure 1.**
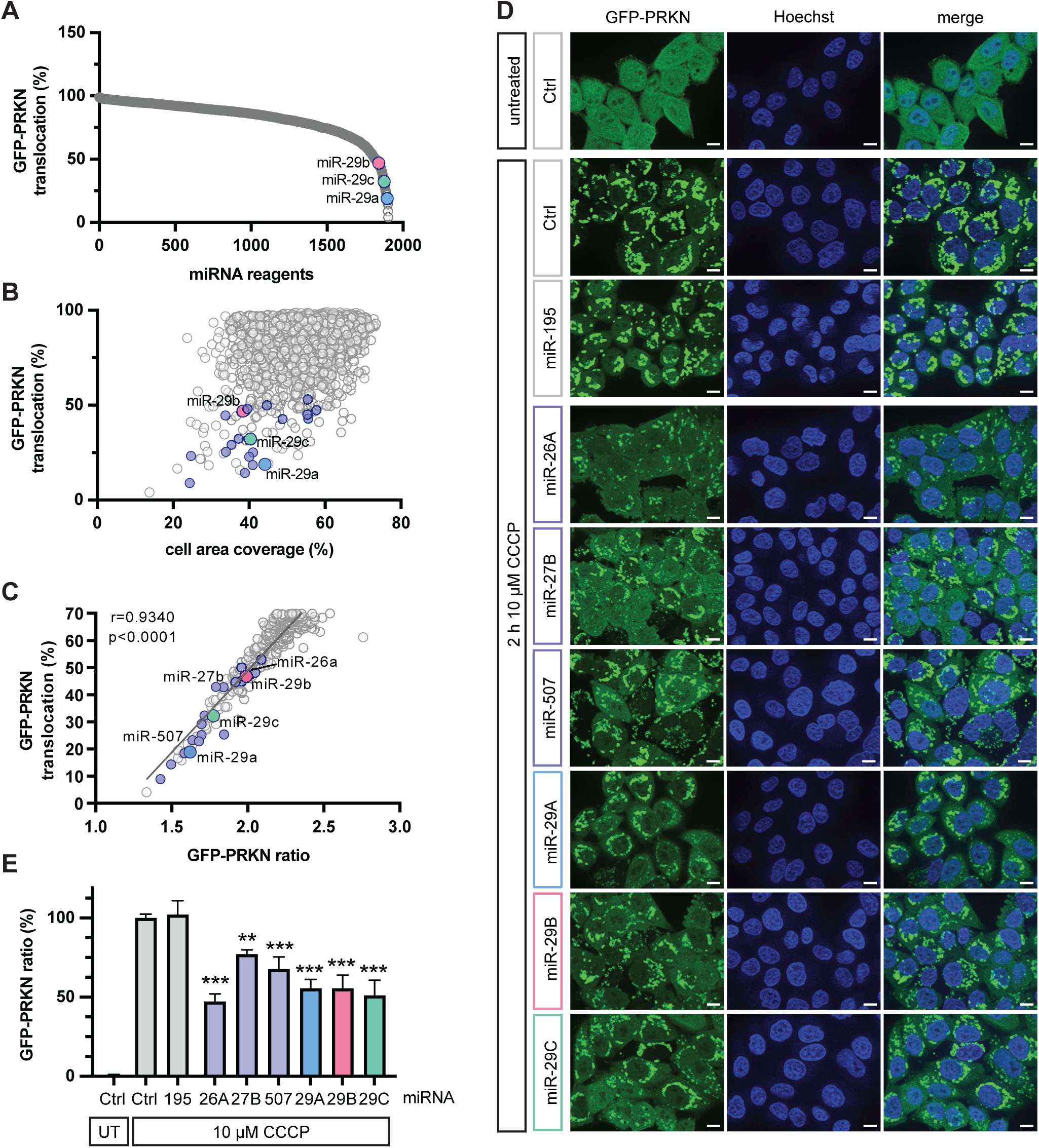
Genome-wide miRNA screening and validation of hits. (**A**) A genome-wide screen with 1902 miRNAs was performed to discover inhibitors of PRKN translocation to damaged mitochondria. The percentage of PRKN translocation positive cells per well was determined with a supervised machine learning linear classifier. Graph shows screening results sorted by effect size. (**B**) PRKN translocation positive cells were graphed against the well percentage covered with cells after miRNA transfection to account for miRNA toxicity to cells. (**C**) Results of the machine learning classifier correlated highly with an established algorithm for PRKN translocation, GFP-PRKN ratio, especially for wells with < 70% PRKN translocation positive cells. (**D, E**) In-house validation of miR-26a, miR-27b, miR-507 and miR-29a/b/c using independent reagents, experimental conditions and a different imaging platform. HeLa cells stably expressing GFP-PRKN were transfected with 0.5 µM miRNA using Lipofectamine RNAiMAX, and treated with 10 µM CCCP for 2 h. A non-specific miRNA (Ctrl) and miR-195 were used as negative controls. (**D**) Representative 63x images are shown. Scale bars correspond to 10 µm. (**E**) Quantification of results using the GFP-PRKN ratio algorithm. Shown is the average GFP-PRKN ratio of 18 wells per experiment ± SEM normalized to the untreated (UT) and treated control transfected cells. Statistical analysis was performed using one-way ANOVA with Dunnett’s multiple comparisons test.

We next randomly selected 20 out of the 86 miRNAs from an independent miRNA library and confirmed them in a follow-up experiment (see **Fig. 1B**, annotated in color). We further selected a few of those candidates with weaker or stronger PRKN translocation effects for additional validation with a higher CCCP concentration and using independent image instrumentation and analysis (**Fig. 1C, D**). Consistent with the results from the screen, miR-195 showed no effect and served as negative control, while miR-27b, which we had described as PRKN translocation inhibitor before [20], showed a significant inhibition. In addition, all new miRNA hits that we tested here, miR-26a, miR-507, and all three members of the miR-29 family, significantly reduced PRKN translocation, suggesting that some of their target genes play a modulatory role in the PINK1-PRKN pathway.

### Identification and prioritization of miR-29a target genes

All three members of the miR-29 family strongly inhibited PRKN translocation, raising the confidence in miR-29 as one of the major miRNAs that influence the PINK1-PRKN pathway. Because of the poor overlap between the results from different miRNA-target prediction algorithms, we decided to use transcriptomics to determine miR-29 target genes experimentally. We transfected HeLa cells either with control miRNA or with miR-29a and performed RNA sequencing. With a stringent cut-off and adjusted p-value of less than 0.01 we received 146 differentially expressed genes (DEGs) (**Fig. 2A**). 122 out of the 146 DEGs (84%) were downregulated, as expected upon overexpression of a miRNA. When overlapping the DEGs with the results from a miRNA target-prediction database, TargetScan, 78 genes overlapped (**Fig. 2B**). For general validation of the RNAseq, we selected several downregulated genes with a broad range of fold changes (**Fig. S2A**) and used quantitative reverse transcription PCR (qRT-PCR) for their assessment. Using independently transfected cell samples, we confirmed all expected gene expression changes including those with lower fold change for each member of the miR-29 family. (**Fig. S2B**).

**Figure 2.**
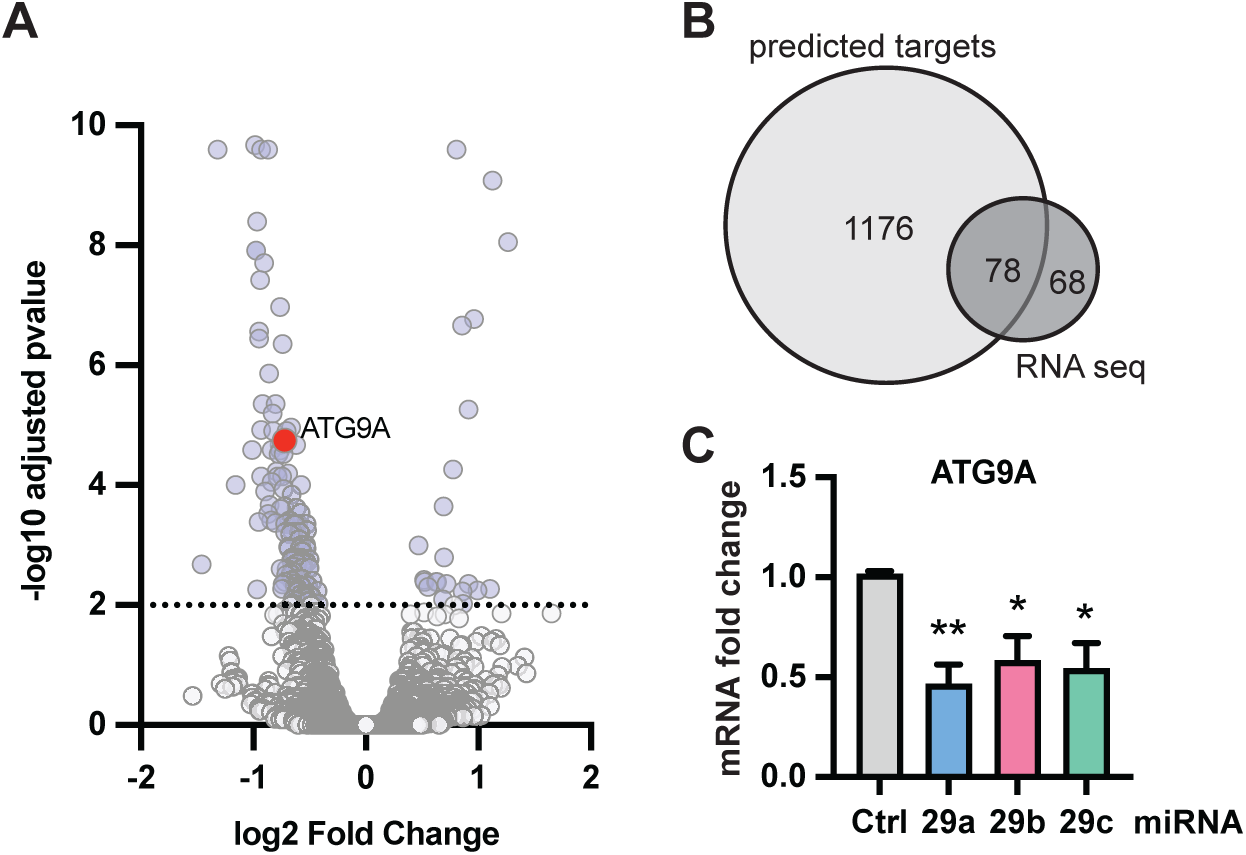
Identification of miR-29a target genes. RNA sequencing of miR-29a was performed as three independent repeats 24 h after transfection of miR-29a. (**A**) Transcriptome data was plotted as volcano plot of fold change and adjusted p-value to visualize results. ATG9A is highlighted in color. (**B**) Of the 146 hits from RNA sequencing, 78 of those were predicted targets of miR-29a according to the TargetScan website [69] (**C**) Cells were transfected with miR-29a, miR-29b, and miR-29c and analyzed for fold change of ATG9A mRNA. Shown is the average fold change ± SEM from at least three independent experiments. Statistical analysis was performed using one-way ANOVA with Dunnett’s multiple comparisons test.

To prioritize miR-29a target genes for mechanistic follow up, we looked for known mitophagy regulators. ATG9A stood out because of its obvious role as core autophagy protein and several reports that connect it to OPTN which plays an important role as adapter for mitophagy [26,27]. ATG9A is a member of the KEGG mitophagy pathway (KEGG ID 04137) and was identified as mitophagy regulator in a previously published CRISPR screen [19]. Furthermore, ATG9A has a miR-29 family target site in the 3’UTR (**Table S4**) and the regulation of ATG9A mRNA was confirmed also by independent qRT-PCR analysis (**Fig. 2C**).

### ATG9A regulates PRKN translocation and is a common target of the miR-29 family

Based on the strong ties of ATG9A with mitophagy, we decided to focus on ATG9A as a candidate gene. We used siRNA to knock down ATG9A and treated siRNA transfected GFP-PRKN cells with CCCP. High content imaging was used to quantify PRKN translocation to damaged mitochondria (**Fig. 3A, B**). PINK1 siRNA, which was included as positive control, led to almost complete suppression of PRKN translocation, as expected. Knockdown of ATG9A significantly decreased PRKN translocation upon CCCP treatment (**Fig. 3B**), similar to miR-29a overexpression.

**Figure 3.**
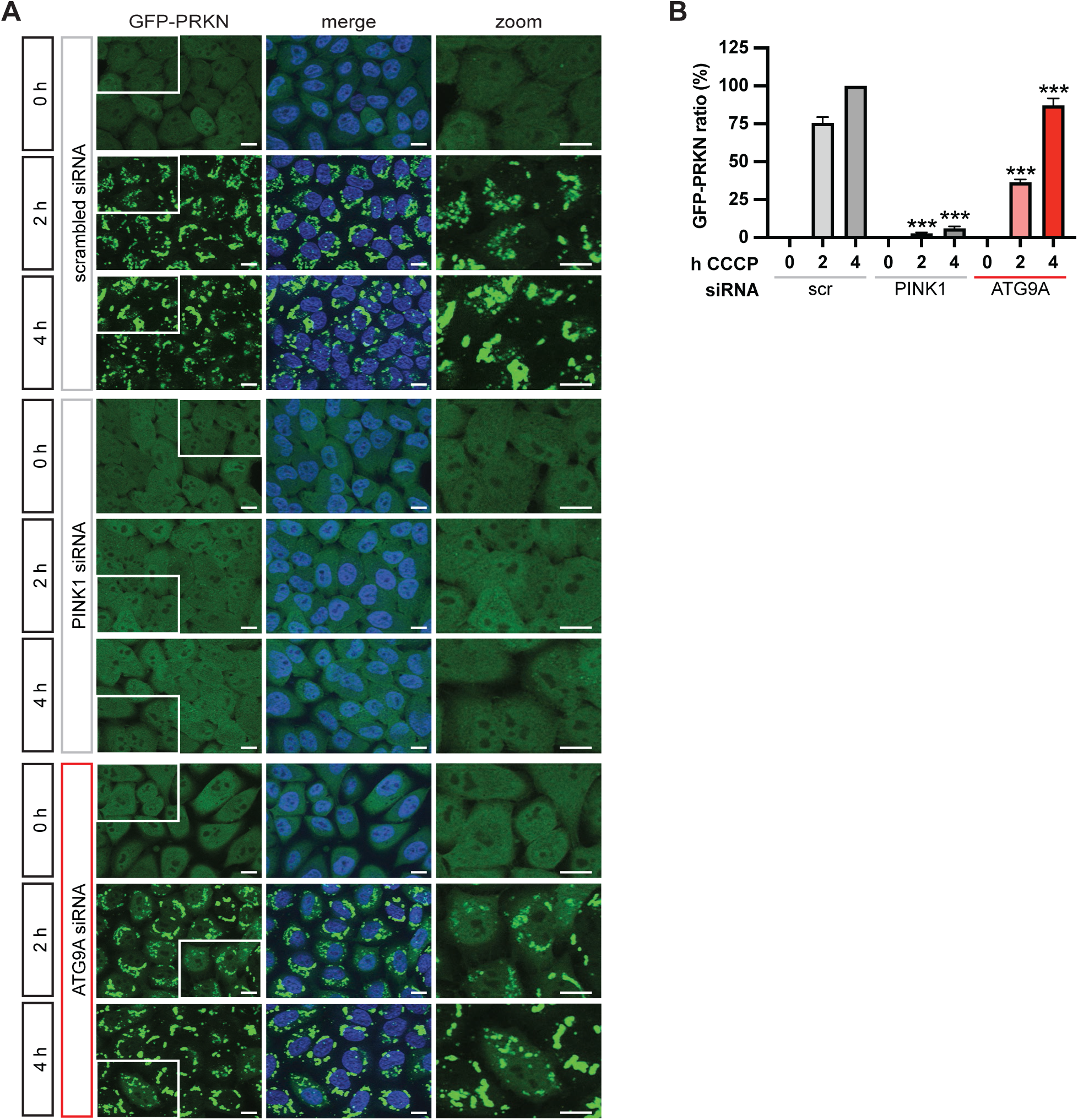
Knockdown of ATG9A inhibits PRKN translocation. (**A**) HeLa GFP-PRKN cells were transfected with scrambled siRNA or siRNA against PINK1 or ATG9A and treated with CCCP for the indicated times. Cells were fixed, and nuclei counterstained with Hoechst. Representative 63x images are shown together with a zoomed in region for each condition. Scale bars correspond to 10 µm. (**C**) Shown is the average GFP-PRKN ratio + SEM of five independent experiments. Data was normalized to values from scrambled siRNA (scr) after 4 h CCCP treatments. Statistical analysis was performed using two-way ANOVA with Dunnett’s multiple comparisons test.

Next, we tested if ATG9A was regulated by miR-29 on protein level. ATG9A protein levels were reduced upon transfection with each individual family member, although this did not reach significance for all conditions (**Fig. 4A, B**). Interestingly, while we initially included PINK1 siRNA as a positive control for reduced mitophagy, we found that samples from cells treated with PINK1 siRNA showed increased ATG9A protein levels (**Fig. 4A, B**), suggesting that ATG9A levels might be upregulated as a response of inhibited mitophagy. This counteracts the miR-29 dependent reduction of ATG9A protein levels upon CCCP treatment. Consistent with a strong effect on PRKN translocation, PINK1 siRNA and all three miR-29 family members led to strongly reduced pS65-Ub levels in CCCP-treated cells (**Fig. 4A, Fig. S3A**). Furthermore, although PINK1 does not have a miR-29 binding site and a meaningful reduction of PINK1 levels was not observed using RNAseq (log2 fold change = -0.261, adjusted p value = 0.64) or by qRT-PCR (**Fig. S3B**), PINK1 protein levels were also significantly decreased upon transfection of all three miRNAs (**Fig. S3C**), suggesting that the inhibitory effect of miR-29 might be amplified via an indirect effect on PINK1.

**Figure 4.**
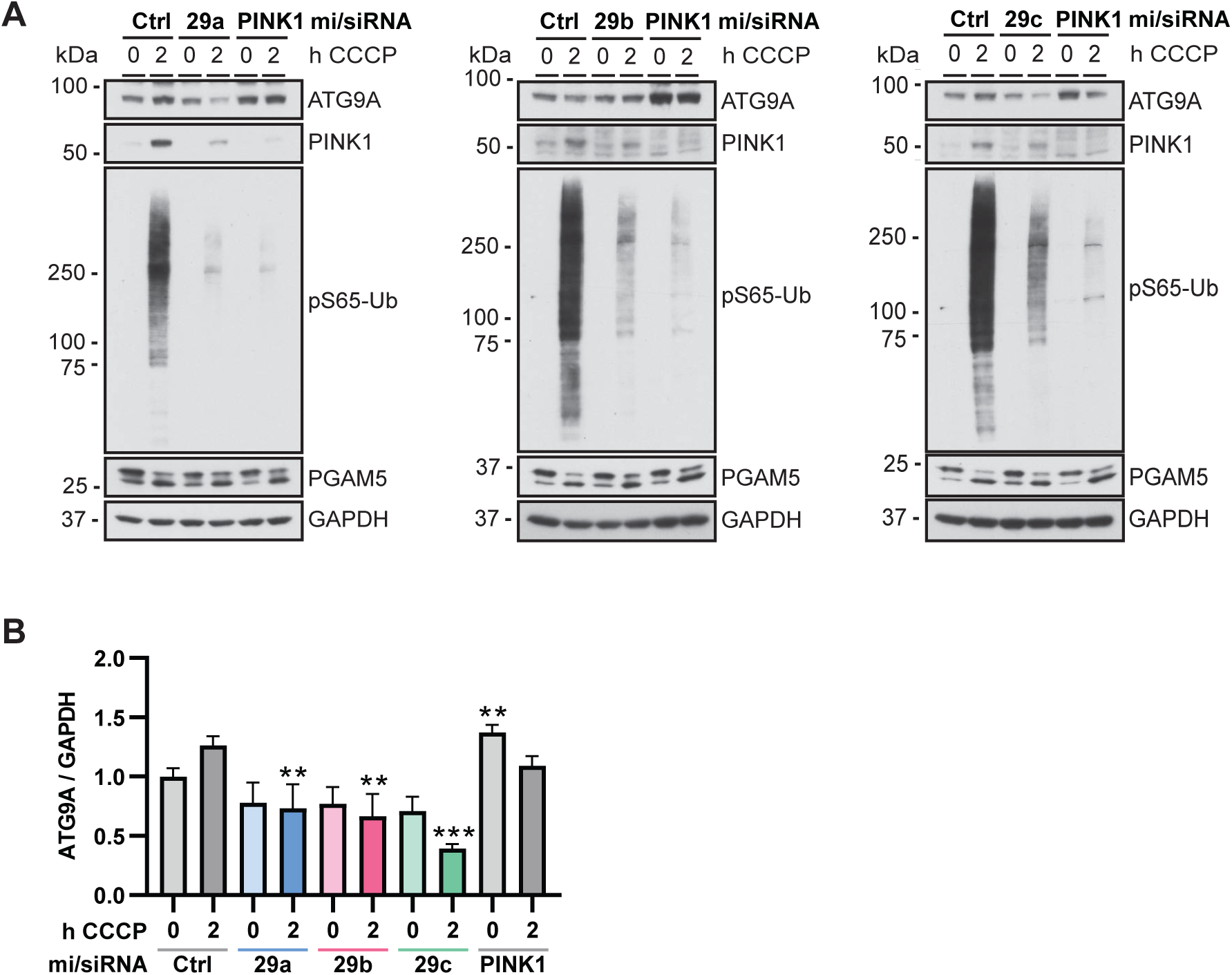
miR-29 expression leads to reduction of ATG9A protein levels. (**A**) Immunoblot analysis of HeLa cells treated with CCCP for the indicated times. Cells were transfected with miR-29a, miR-29b, or miR-29c and PINK1 siRNA as a control and analyzed via Western blot for effects on ATG9A, PINK1, and pS65-Ub. PGAM5 was used as control for the successful depolarization of the cells, GAPDH as loading control. (**B**) Shown is the mean ATG9A levels normalized to GAPDH ± SEM of at least three independent experiments. Statistical analysis was performed using two-way ANOVA with Dunnett’s multiple comparisons test.

### Knockdown of ATG9A inhibits mitophagy and expression rescues miR-29 mediated mitophagy inhibition

Knockdown of ATG9A with siRNA significantly reduced its own mRNA (**Fig. 5A**) and protein levels (**Fig. 5B, C**). Transfection of cells with siRNA against ATG9A further led to a small but significant decrease of PINK1 mRNA levels (**Fig. 5D**). At 6 h after CCCP treatment, PINK1 protein levels also showed a trend toward reduction upon ATG9A siRNA but this was not significant (**Fig. 5B, C**). As a readout of PINK1-PRKN activation, we monitored pS65-Ub by western blot (**Fig. 5B**) and quantified its levels using a sandwich ELISA (**Fig. 5E**). In line with the high content imaging results, knockdown of ATG9A resulted in significantly decreased pS65-Ub levels. We further assessed mitophagy using the mitoKeima reporter, a mitochondrial targeted, pH-responsive probe that can be used to measure the acidification that occurs when mitochondria are delivered to lysosomes for degradation [28]. HeLa GFP-PRKN cells stably expressing mitoKeima were transfected with scrambled siRNA, ATG9A siRNA, or PINK1 siRNA as positive control and live imaged for 18 h after CCCP treatment to monitor the signal of neutral and acidic mitoKeima (**Fig. S4A**). Compared to control cells, ATG9A siRNA significantly reduced the total area of acidic mitochondria over total mitochondria per cell starting at the 6 h CCCP treatment, corroborating that knockdown of ATG9A inhibits mitophagy (**Fig. S4B**).

**Figure 5.**
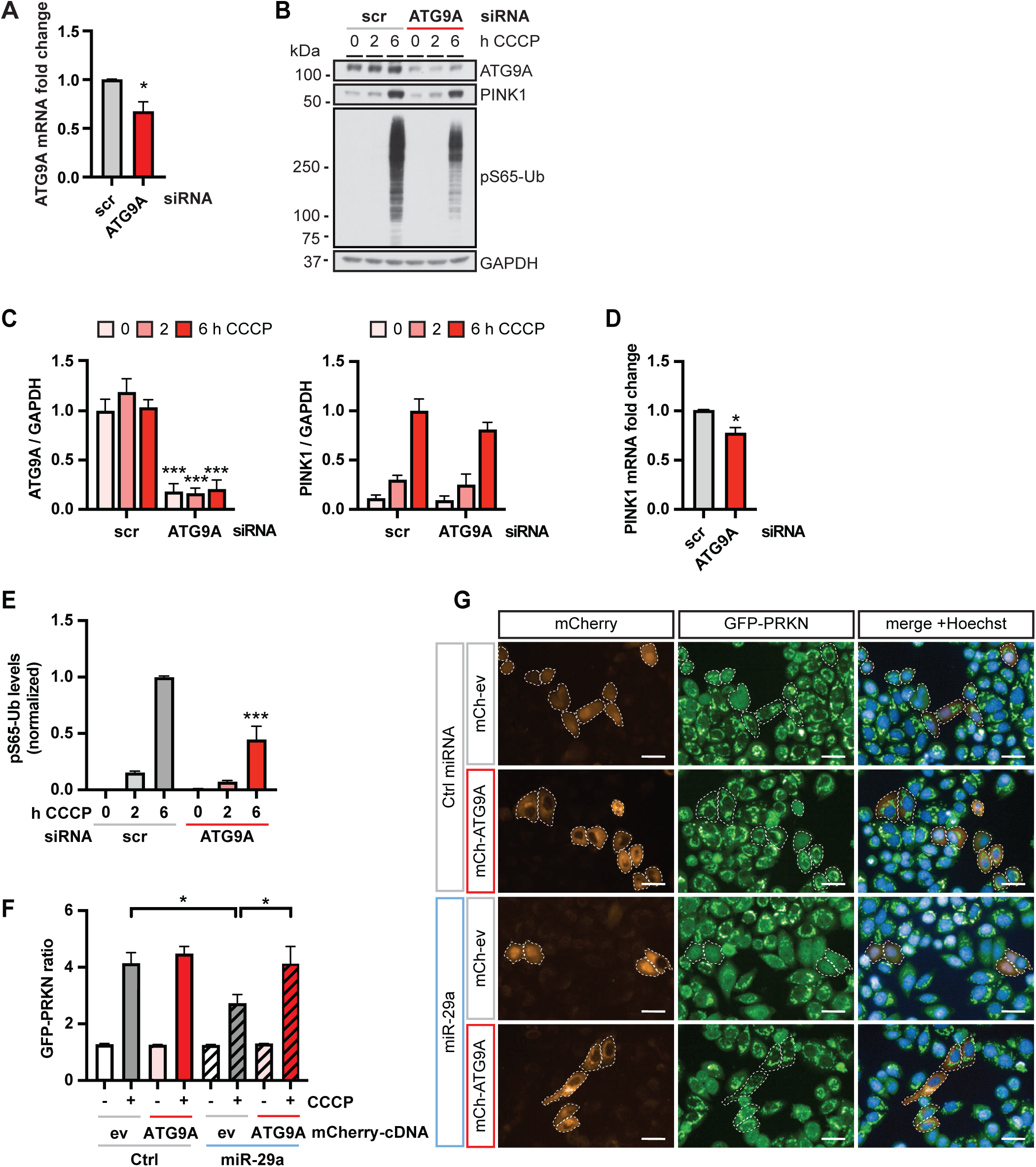
Knockdown of ATG9A inhibits and re-expression restores mitophagy readouts. (**A**) HeLa cells were transfected with 20 µM siRNA to silence ATG9A and analyzed for fold change of mRNA levels via qRT-PCR. Shown is the average fold change normalized to the scrambled control (scr) + SEM. Statistical analysis was performed using an unpaired, two-sided t-test. (**B**) Cells were transfected with scrambled (scr) or ATG9A siRNA and analyzed via Western blot for effects on the target gene and PINK1. (**C**) Protein levels were quantified and normalized to GAPDH as loading control. Shown is the normalized mean + SEM of three independent experiments. Statistical analysis was performed using two-way ANOVA with Sidak’s multiple comparisons test. (**D**) HeLa cells were transfected with 20 µM ATG9A siRNA and analyzed for fold change of mRNA levels of ATG9A via qRT-PCR. Shown is the average fold change normalized to the scrambled control (scr) + SEM. Statistical analysis was performed using an unpaired, two-sided t-test. (**E**) Cells were analyzed for pS65-Ub levels using MSD ELISA. Shown is the mean normalized to the scrambled siRNA control + SEM of at least three independent experiments. Statistical analysis was performed using two-way ANOVA with Dunnett’s multiple comparisons test. (**F**) GFP-PRKN HeLa cells were transfected with either a negative control miRNA or miR-29a then subsequently transfected with either an empty vector expressing mCherry (mCh-ev) or mCherry-ATG9A and treated with CCCP for the indicated times. Shown are representative images for each condition. Scale bars represent 10 µm. (**G**) Shown is the mean GFP-PRKN ratio + SEM from four independent experiments. Only mCherry positive cells with intensity >1500 were analyzed using the GFP-PRKN ratio algorithm. Statistical analysis was performed using two-way ANOVA with Dunnett’s multiple comparisons test.

In order to determine whether ATG9A was indeed primarily responsible for the inhibitory effect of miR-29, we performed rescue experiments. We transfected cells either with a control miRNA or with miR-29a in combination with an empty mCherry vector or mCherry-tagged ATG9A and then performed high content imaging where we analyzed GFP-PRKN translocation only in mCherry-positive cells (**Fig. 5F, G**). Transfection of miR-29 effectively reduced the GFP-PRKN ratio to about 60% in empty vector transfected cells, while co-transfection of ATG9A fully restored GFP-PRKN translocation in these cells (**Fig. 5F, G**). Together our functional analysis suggests that ATG9A is a key effector of miR-29a since its knockdown reduced mitophagy and its re-expression restored GFP-PRKN translocation in miR-29a transfected cells.

In order to shed light onto the potential mechanism by which silencing of the downstream autophagy effector ATG9A would inhibit PRKN translocation upstream, we stained cells with a BODIPY dye to visualize lipid droplets. A recent paper discovered that ATG9A functions as a lipid scramblase that facilitates mobilization from lipid droplets to mitochondria, which in turn enables mitochondrial fatty acid β-oxidation [29]. Consistent with this, RNA interference of ATG9A resulted in significantly more lipid droplets and a higher lipid droplet area per cell compared to the controls in HeLa cells (**Fig. S5A, B**), although the effect was weaker compared to the original study where ATG9A was knocked out [29]. Interestingly, transfection with miR-29 also led to increased lipid droplet number and area (**Fig. S5C, D**). In neither case did the lipid droplet number and size change upon CCCP treatment. In line with this, CCCP treatment also did not seem to induce any alterations of ATG9A protein localization (**Fig. S5E**). ATG9A, which is known to be localized to Golgi vesicles [30] did not overtly co-localize with mitochondria.

### EOPD-associated ATG9A variants fail to restore miR-29a mediated mitophagy inhibition

We next explored an EOPD cohort consisting of 208 individuals with an age at onset before age of 50 years for mutations in *ATG9A*. Using available whole genome sequencing data, we focused on rare, exonic variants and found two samples each with one non-synonymous heterozygous variant. Both resulting substitutions, ATG9A p.R631W and p.S828L, had a much lower allele frequency in the gnomAD reference database (**Fig. 6A**). The ATG9A p.R631W variant was found in a white male patient who was diagnosed with PD at 43 years of age and responsive to L-DOPA. Initial symptoms included left hand tremor and gait difficulties. The p.S828L variant was identified in a female patient with Hispanic or Latino ethnicity, who developed symptoms at 43 years of age. Onset symptoms included asymmetry, bradykinesia, gait difficulties, rigidity and resting tremor. This individual was unresponsive to anti-parkinson therapy. Results of combined annotation dependent depletion (CADD) analysis for ATG9A p.R631W and p.S828L suggested that both variants are among the most 1% deleterious in the human genome (**Fig. 6A**). Both variants lead to changes of highly conserved amino acids in the C-terminal intrinsically disordered domain of ATG9A outside of the transmembrane core (**Fig. 6B, Table S5**). Of note, when testing these variants experimentally in miR-29a rescue experiments, both mutations failed to restore PRKN translocation and resulted in significantly reduced PRKN translocation compared to ATG9A wildtype (**Fig. 6C, D**). This suggests that these variants act as loss-of-function variants and opens up the possibility that they might be related to the etiology of PD.

**Figure 6.**
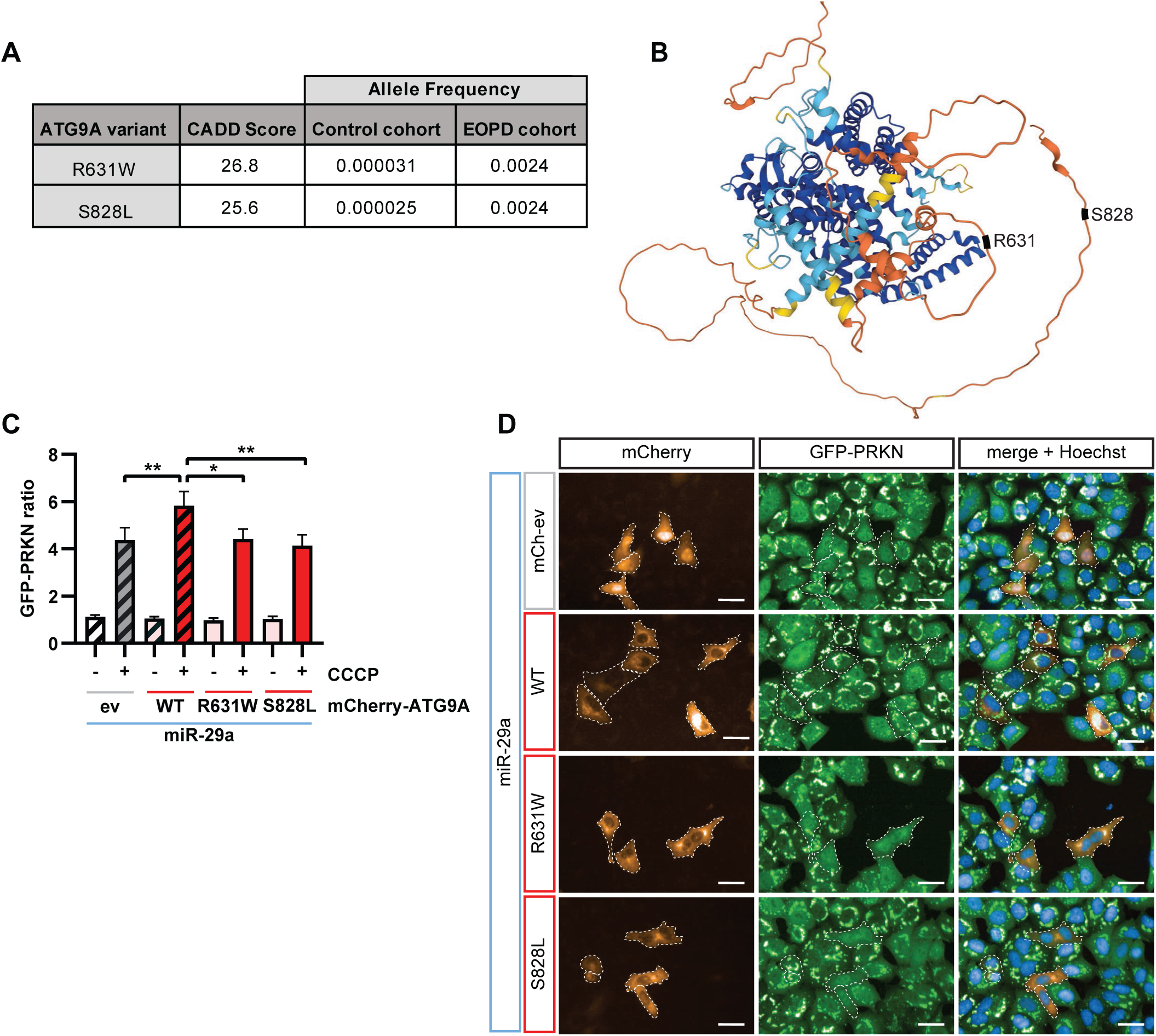
EOPD associated ATG9A variants lead to loss of function. (**A**) Shown are the CADD scores, frequencies of the control cohort from genome database gnomAD [70], and the frequencies from our EOPD cohort for the two *ATG9A* variants p.R631W and p.S828L. (**B**) Shown is the predicted protein structure of ATG9A, highlighting the locations of the EOPD-associated variants in disordered regions of the protein. Structure prediction obtained from AlphaFold [71]. (**C**) GFP-PRKN HeLa cells were transfected with miR-29a and then subsequently transfected with an empty mCherry vector (ev), mCherry-ATG9A wildtype (WT), R631W or S828L and treated for the indicated times. Only mCherry positive cells with intensity >1500 were analyzed using the GFP-PRKN ratio algorithm. Shown is the mean + SEM from three independent experiments. Statistical analysis was performed using two-way ANOVA with Dunnett’s multiple comparisons test. (**D**) Shown are representative images for each condition. mCherry-positive are outlined with a white dashed line for illustrative purposes. Scale bars represent 10 µm.

## DISCUSSION

Using a genome-wide high content imaging miRNA screen, we have discovered novel regulators of PRKN translocation and thus mitophagy. Of the 86 inhibiting miRNAs, we validated 20 using independent chemistry and, because all three members significantly reduced PRKN translocation, selected the miR-29 family for functional follow-up. We further experimentally determined miR-29 target genes by RNAseq (n=146) and selected ATG9A for validation. siRNA-mediated downregulation of ATG9A phenocopied the effects of miR-29 and significantly inhibited PRKN translocation. Importantly, re-expression of wild-type ATG9A, but neither of two rare and potentially deleterious ATG9A substitutions identified in EOPD patients was able to restore PRKN translocation. Together, our study identifies miR-29 as a new mitophagy regulator and shows that it is mediated via ATG9A. Our study further supports a potential role for the loss-of-function of other PINK1-PRKN mitophagy proteins, such as ATG9A, for the etiology of EOPD.

As the most upstream regulator of this pathway, PINK1 is a likely target of the inhibitory miRNAs we identified. However, of the five miRNAs that have been experimentally confirmed to directly target PINK1 [31–33], only miR-27b [20] was among the top hits in our genome-wide screen. Other miRNAs that have been indirectly linked to PINK1 signaling before, such as miR-155, which regulates PINK1 stability through BAG5 [34], miR-330 and miR-21-5p, which both regulate the mitochondrial phosphatase PGAM5 [35,36], or miR-421, which is a predicted regulator of PINK1 that was shown to inhibit mitophagy, were also not uncovered in our screen. While PINK1 is neither a predicted or experimental target of miR-29, we cannot rule out that direct or indirect changes in PINK1 levels caused some of the observed effects in our study. In fact, transfection with miR-29 or siRNA against ATG9A caused reduction of PINK1 on mRNA and/or protein level, which might exacerbate mitophagy inhibition. However, the fact that re-expression of ATG9A fully recovered the PRKN translocation deficits that were induced by miR-29 suggests that ATG9A plays a critical role for this process.

Several studies have identified miRNAs dysregulated in PD, but the lack of overlap between the reports that were mostly limited by a small sample size has led to an ambiguous picture [37]. Nevertheless, miR-29 and five other miRNAs (miR-26 and miR-30, miR-199b, miR-212, miR-380-5p) out of 89 miRNAs that have been found to be upregulated in sporadic human PD brain [38–40] inhibited PRKN translocation by at least 50% in our screen. In peripheral PD samples such as blood and CSF, miR-29 was mostly found reduced with discordant results among studies and differences among individual miR-29 family members (reviewed in [41]). In our screen, all three members of the miR-29 family inhibited PRKN translocation. miR-29a, which is the most abundantly expressed and stable among the family [42], had the strongest effect, followed by miR-29c and miR-29b. miR-29b and miR-29c have a tri-uracil element at nucleotide positions 9-11 that causes their rapid decay [42]. All members share the same seed sequence and therefore have similar targets. miR-29a is mostly cytosolic, while miR-29b and miR-29c have been found in the nuclear fraction of HeLa cell lysates [43]. Besides PD, the miR-29 family has been linked to other diseases, such as leukemia, osteoporosis, and various heart diseases [44–46].

siRNA-mediated silencing of ATG9A resulted in reduced PRKN translocation to damaged mitochondria and lower pS65-Ub levels. ATG9A is a core autophagy transmembrane protein that is packaged in vesicles formed from the Golgi. ATG9A, which itself is not incorporated into autophagosomes, plays an essential role for their formation and expansion by functioning as a lipid scramblase [47]. Together with IQGAP1 and ESCRT proteins, ATG9A has also been shown to protect against plasma membrane damage [48]. Both full and conditional brain ATG9A KO mice die a very early premature death [49,50]. The observed accumulation of p62/SQSTM1, NBR1 and ubiquitin suggest that autophagy is inhibited in ATG9A KO animals [49,50]. In line with this, ATG9A knockout in HeLa cells and mouse embryonic fibroblasts results in fewer, but larger LC3 puncta that have been shown to colocalize with ubiquitinated p62 structures [51,52]. ATG9A has been previously reported to interact with other important mitophagy proteins, including RAB7A, ULK1, AMBRA1, and OPTN [26,53–56]. RAB7A recruits ATG9A to the mitophagosome formation site to promote expansion [56]. ULK1 and ATG9A independently translocate to the autophagosome formation site where they recruit other downstream ATG proteins [54]. While it remains unclear how the autophagosome regulator ATG9A regulates PRKN translocation as one of the earlier steps in the pathway, we found that knockdown of ATG9A and overexpression of miR-29 both altered the number and size of lipid droplets, indicating that a previously identified lipid scramblase function that affects mitochondrial fatty acid β-oxidation [29] might be involved in this process. Interestingly, a role for lipogenesis in PRKN translocation and PINK1 stabilization has been suggested before by the identification of the SREBF1/2 transcription factor in a previous genome-wide siRNA screen [16]. While ATG9A was found among the SREBF1 target genes in ChIP-seq experiments [57], it remains to be studied if there is a mechanistic connection between these two with regards to mitophagy.

Using whole genome sequencing data from an EOPD cohort, we identified two rare nonsynonymous variants in *ATG9A* that we tested functionally. In contrast to re-expression of WT, both ATG9A p.R631W and p.S828L failed to restore PRKN translocation upon miR-29a transfection, suggesting that they act as loss-of-function mutations. This implies that mitophagy deficits might be linked to disease in the carriers. However, if and how heterozygous *ATG9A* variants are really linked to the pathogenesis of EOPD remains to be determined and further genetic studies are warranted. ATG9A forms a homotrimer with a wide, solvated central pore and a lateral cavity [52]. It is conceivable that a heterozygous mutation could disrupt the function of the entire trimeric complex. In fact, we found that inactive ATG9A with mutations in the core part of the protein increased LC3 puncta size not only in ATG9A KO but also in wildtype HeLa cells, suggesting a dominant-negative effect [58]. Alternatively, *ATG9A* haploinsufficiency together with other genetic variants might cause a combined reduction of mitophagy below a critical threshold. Indeed, the individual with the *ATG9A* p.S828L also carries a heterozygous *PRKN* p.R275W mutation. Regardless of the mechanism, a recent study found that ATG9A was reduced in fibroblasts from late-onset PD patients on the translational level [59]. While it is unclear if this is connected to increased expression of miR-29 in these samples, it reiterates a potential role of ATG9A-mediated dysfunctional mitophagy for PD and makes ATG9A an attractive topic for future translational studies. In conclusion, our study provides critical proof of concept and highlights the power of functional cell biological screening to unravel the genetic architecture of mitophagy and EOPD.

## MATERIAL AND METHODS

### Cell Culture

HeLa cells, HeLa cells stably overexpressing GFP-PRKN [21] alone or with mitoKeima overexpression [24], were cultured in Dulbecco’s modified Eagle’s medium (DMEM, Thermo Fisher Scientific, 11965118) supplemented with 10% fetal bovine serum (FBS, Neuromics, FBS001800112) by volume. All cells were grown at 37°C, 5% CO2/air in humidified atmosphere. For miRNA and siRNA transfection in 12- or 6-well plates, cells were plated 24 h before transfection. To transfect a 6-well plate with miRNA, 18 µl of RNAiMAX (Thermo Fisher, 13778075) was added to 150 µl Opti-MEM (Thermo Fisher Scientific, 51985-034) and 6 µl of 10 µM miRNA was added to 150 µl Opti-MEM. Once mixed, 250 µl of mixture was added dropwise to the cells, for a final miRNA concentration of 25 nM. For siRNA transfection in a 12-well plate, 50 µl Opti-MEM, 6 µl Hi-Perfect transfection reagent (Qiagen, 301707), and 1 µl of the desired (20 µM) siRNA were mixed. The mixture was added dropwise to the cells for a final siRNA concentration of 10 nM. The siRNAs used were ATG9A (Horizon Discovery, D-014294-02-0010), PINK1 (Qiagen, SI00287931), and AllStars Negative control (Qiagen, 1027281). The following miRNAs from Qiagen were used: miRNA29a (SI05352109), miRNA29b (SI05352501), miRNA29c (SI05358486), miRNA195 (SI05356190), miRNA27b (SI05354874), miRNA26a (SI05351997), miRNA507 (SI05364107), and AllStars Negative control (SI03650318). Carbonyl cyanide m-chlorophenyl hydrazone (CCCP) (Sigma-Aldrich, C2759-100MG) was used as treatment.

### Genome-wide miRNA screen and validation

The genome-wide miRNA screen was performed at the Sanford Burnham Prebys Medical Discovery Institute with 1902 miRNAs (MISSION version 17 library, 1902 miRNAs, Sigma-Aldrich, MI00200). Briefly, 650 cells per well were reverse transfected in 384-well plates (Greiner, 781092) using 10 nM of miRNA with Lipofectamine RNAimax. Cells were incubated for 72 h before they were treated with 3.5 µM CCCP for 2 h. Cells were fixed by adding a 16% PFA solution (Electron Microscopy Sciences, 15710) for 30 min at room temperature for a final concentration of 5% PFA. Plates were then washed three times with PBS (Invitrogen, 14190-235) and then left in PBS with Hoechst (Invitrogen, H21492, 1:4000 final concentration) for at least 30 min before imaging. Controls included the library internal negative controls ath-miR416 and cel-miR-243 as well as pooled non-specific siRNAs [60,61] that were transfected into cells and then either treated with DMSO (Sigma, D4540-500ML), with 0.7 µM CCCP, or with 3.5 µM CCCP to mimic absent, weak or strong PRKN translocation, respectively. A killer siRNA (eIF4A3, Ambion) [22] was used to estimate the transfection efficiency in comparison to some wells that were not transfected or transfected with the non-specific siRNAs. Plates were imaged on a PerkinElmer Opera QEHS (PerkinElmer, Waltham, MA, USA) with a 20x magnification using a 488nm laser with a 540/75 emission filter and xenon arc lamp excitation at 365 nm with a 450/40 nm emission filter. Each miRNA was tested in duplicate, and four images were taken per well.

Image analysis was performed in Signals Image Artist, version 1.0 (PerkinElmer, Waltham, MA). In brief, nuclei and cell borders were identified. A population of cells was selected based on size, roundness, and without contacting the image border. 415 standard morphology, advanced morphology, and intensity properties were considered by the supervised machine learning linear classifier (PhenoLOGIC) to classify cells into two main groups: cells with diffuse PRKN (DMSO-treated wells), cells with translocated PRKN (3.5 µM CCCP treated wells). The algorithm further identified a third, “InBetween” phenotype that had characteristics of both positive and negative controls. For the purpose of the screen the cells with this phenotype were considered PRKN translocation negative. As alternative readout for PRKN translocation, the ratio of cytoplasmic maximal intensity to nuclear mean intensity of GFP was used. For validation, miRNAs were picked from an independent library (Applied Biosystems, AM17105), which included 875 human miRNAs listed in miRBase sequence database version 10.1.

### In-House High Content Imaging

For in-house high content imaging, reverse transfection of siRNA and miRNA was performed in 96-well (PerkinElmer, 6055308) or 384-well (Greiner, 781092) plates. For siRNA transfection, 500 µl Opti-MEM, 7.5 µl Hi-Perfect transfection reagent (Qiagen, 301707), 1.25 µl siRNA stock (20 µM) was mixed and 10 µl were added per well on a 384-well plate. HeLa pWPI EGFP-Myc-PRKN cells were plated on the transfection mixture at 1000 cells per well. For reverse transfection of miRNA, 2 µl 0.5 µM miRNA and 10 µl 1:50 Lipofectamine RNAimax were added per well on a 384-well plate. HeLa GFP-PRKN cells were plated on the transfection mixture at 1400 cells per well. 72 hours after transfection, cells were treated with 20 µM CCCP. Cells were fixed for 10 min with 4% PFA and stained with Hoechst 33342 (1:5000). Plates were imaged using Operetta CLS High Content imaging system (Perkin Elmer, Waltham, MA, USA) equipped with Harmony analysis software. For Harmony-based PRKN translocation analysis, the ratio of the maximum GFP intensity in the cytoplasm and the mean GFP intensity in the nucleus was used. For rescue experiments, GFP-PRKN ratio was only analyzed in mCherry-positive cells with an intensity between 1500 and 4000.

For lipid droplet analysis, a binary image was engineered by first smoothing and removing the background of the image using a sliding parabola curvature of 20. Any pixels with an intensity greater than 800 were kept in the image. Spots within cells were identified using the “Find Spots” building block, keeping the detection, and splitting sensitivities low.

In order to analyze mitophagic flux, GFP-PRKN cells that stably also expressed mitoKeima [24] were imaged live over time one hour after cells had been incubated with Hoechst. Images were taken every two hours for the first three measurements and every four hours for the remainder of the experiment for a total of 18 hours. For mitophagy flux analysis, we calculated the total area of acidic over total mitochondria per cell.

### Real-time Quantitative PCR

RNA was isolated using the RNeasy kit (Qiagen, 74104) according to the manufacturer’s instructions. RNA concentration was determined using a Nanodrop (Thermo Fisher, Waltham, MA, USA). The cDNA was prepared using the Transcriptor High Fidelity cDNA Synthesis kit (Roche). 1000 ng of RNA was used for reverse transcription in a 6.5 µl reaction using the iTaq Universal SuperMix Two-step kit (Biorad, 1725130). 3 µl of the master mix for each probe and 2ul of the generated cDNA was pipetted into a 384-well plate. mRNA levels were measured by quantitative PCR by using a LightCycler480 (Roche, Rotkreuz, Switzerland) using specific Taqman probes. Relative expression was determined by using RPL27 as a housekeeping gene. The probes were purchased from Bio-Rad and had the following Assay ID: RPL27 (qHsaCEP0051648), ATG9A (qHsaCEP0055662), PINK1 (qHsaCEP0053094), COL4A1 (qHsaCIP0027494), HMGCR (qHsaCEP0050733), CDC42 (qHsaCIP0040659), and VEGFA (qHsaCEP0050716). Relative quantification was performed using the ddCT method [62].

### RNAseq and data analysis

RNA concentration and quality were initially assessed using Qubit fluorometry (ThermoFisher Scientific, Waltham, MA) and the Agilent Fragment Analyzer (Santa Clara, CA). cDNA libraries were prepared using 200 ng of total RNA according to the manufacturer’s instructions for the TruSeq Stranded mRNA Sample Prep Kit (Illumina, San Diego, CA). The concentration and size distribution of the completed libraries were determined using Qubit fluorometry and an Agilent BioAnalyzer DNA 1000 chip (Santa Clara, CA). Libraries were multiplexed into a single sequencing lane following Illumina’s standard protocol using the Illumina cBot and HiSeq 3000/4000 PE Cluster Kit. The flow cell was sequenced as 100 X 2 paired end reads on an Illumina HiSeq 4000 using HiSeq 3000/4000 sequencing kit and HD 3.4.0.38 collection software. Base-calling was performed using Illumina’s RTA version 2.7.7. Paired-end sequencing was performed on mRNA collected from HeLa cells of 6 total samples, with an n=3, for each, control miRNA and miR-29a transfected samples. The secondary analysis of the sequenced samples was performed using the Mayo Clinic developed workflow MAP-Rseq version 3.0.0 [63]. This pipeline implements the alignment of paired end reads to the hg38 reference genome using STAR [64] aligner. Gene and expression level quantification were derived using Subread [65] to calculate both the raw and normalized Reads Per Kilobase per Million (RPKM) values. For overall quality control analysis of the sequenced libraries, the RSEQC [66] package is run on the aligned reads. Differential Expression Analysis was performed using edgeR [67]. Prioritized results were determined using an adjusted p value < 0.01.

### Western Blot

Cells were lysed in RIPA buffer (50 mM Tris [Sigma-Aldrich, 648311], pH 8.0, 150 mM NaCl [Sigma-Aldrich, S5886], 0.1% SDS [Fisher Scientific, BP166-500], 0.5% Deoxycholate [Sigma-Aldrich, D6750], 1% NP-40 [Sigma-Aldrich, I3021]) with addition of Complete protease and PhosSTOP phosphatase inhibitor (Sigma, 11697498001, 4906837001). A bicinchoninic acid assay (Fisher Scientific, PI-23225) was used to determine protein concentration. Protein samples were boiled at 95°C, aside from ATG9A probed samples, and transferred onto polyvinylidene fluoride (PVDF) membranes (Millipore, IEVH00005). Membranes were blocked for one hour at room temperature in 5% dry milk (Sysco, 5398953) dissolved in TBST (500 mM Tris, 1.5 M NaCl, 0.1% Tween20 [Sigma, 9005-64-5]) followed by overnight incubation with primary antibodies at 4°C. Membranes were incubated with HRP-conjugated secondary antibodies (1:10,000; Jackson Immunoresearch Laboratories) in 5% milk in TBST for one hour at room temperature. Bands were visualized with Immobilon Western Chemiluminescent HRP Substrate (Millipore, WBKLS0500) on Prometheus ProSignal ECL Blotting Films (Genesee, 30-810L) or by using ChemiDoc MP Imaging System (Biorad, Hercules, CA, USA). Western blot analysis was performed by using Image Studio Lite software version 5.2.5.

### Immunofluorescence Staining

Cover glass slips were coated with Poly-D-lysine hydrobromide (Sigma-Aldrich, P6407). Cells were plated on the glass coverslips in 24-well plates. Cells were washed with 1x PBS (Invitrogen, 14190-235), fixed with 4% PFA (Fisher Scientific, AAJ19943K2) in PBS for 10 minutes, and washed again with PBS. Cells were then blocked for one hour with 10% goat serum (Invitrogen, 16210072) in PBS. Primary and secondary antibodies were diluted in 1% BSA in PBS and incubated for one hour each. Cells were fixed once more with 4% PFA in PBS for 10 minutes. For lipid droplet analysis, cells were washed once with PBS supplemented with 0.1 mM CaCl2 and 1.0 mM MgCl2 (PBSCM). Cells were fixed with 4% PFA in PBSCM and then washed once with PBSCM. Cells were permeabilized with PBSCM with 0.1% saponin (Sigma-Aldrich, S7900) and 0.1% BSA for 10 min at room temperature. Cells were stained with BODIPY 493/503 (Fisher Scientific, D3922) for 30 min at 37°C (1 µg/ml in PBSCM containing 0.1% saponin and 0.1% BSA, 30ul per well) with Hoechst 33342 (1:1000). Representative images were obtained by AxioObserver microscope equipped with an ApoTome Imaging System (Zeiss, Oberkochen, Germany).

### Antibodies

The following primary antibodies were used for western blot analysis: rabbit anti-ATG9A (Abcam, ab108338, 1:1000), mouse anti-GAPDH (Meridian Life Sciences, H86504M, 1:750,000), mouse IgG2b anti-PINK1 (Novus, NBP2-36488, 1:10,000), rabbit anti-phospho S65 ubiquitin (Cell Signaling Technologies, 62802, 1:20,000), rabbit anti-PGAM5 (Abcam, AB126534, 1:15,000). The following secondary antibodies from Jackson Immunoresearch were used for western blot analysis: Peroxidase-AffiniPure donkey anti-mouse IgG (H+L) (715-035-150), Peroxidase-AffiniPure donkey anti-rabbit IgG (711-035-152), Peroxidase-AffiniPure goat anti-mouse IgG Fcγ subclass 2b specific (115-035-207). The following primary antibodies were used for immunofluorescence analysis: rabbit anti-ATG9A (Abcam, ab108338, 1:200), mouse IgG1 anti-HSP60 (PTG, 66041-1-Ig, 1:4000). The following secondary antibodies were used for immunofluorescence analysis: goat anti-rabbit AlexaFluor 568 (Molecular Probes, A11011, 1:1000), goat anti-mouse AlexaFluor 647 (Molecular Probes, A21235, 1:1000).

### ELISA

Our in-house Meso Scale Discovery ELISA against p-S65-Ub was published previously [68]. In brief, 96-well Meso Scale Discovery assay plates (MSD, L15XA-6) were coated with 30 μl of pS65-Ub capturing antibody (Cell Signaling Technology, 62802) with a final concentration of 1 µg/ml in 200 mM sodium carbonate coating buffer (pH 9.7) overnight at 4°C. Plates were washed three times with 0.1% Tween-20 in TBS (TBST), blocked in 1% BSA/TBST for 1 h at room temperature and washed again. Protein lysates were prepared in RIPA buffer diluted in 1% BSA/TBST to final concentration 0.5 µg/µl and 30 µl of diluted lysates (total amount: 15 µg of protein lysate) were incubated overnight on the plate at 4°C. Plates were washed and incubated with 30 μl mouse anti-total-Ub detection antibody (1:500, P4D1, Thermo Fisher, 13-6078-82) for 1 h, followed by three washes and incubation with goat anti-mouse SULFO TAG secondary antibody (MSD, R32AC-5, 1:500 in 1% BSA/TBST) for 1h at room temperature. Plates were washed and signal measured after addition of 150 μl/well Gold Read Buffer (MSD, R92TG-2) on a SECTOR Imager 2400 (MSD, Rockville, Maryland, USA) using the MSD workbench Software.

### Genetic screening cohort

The primary genetic analysis for *ATG9A* included 150 unrelated EOPD samples collected at the Mayo Clinic. The average age at onset was 39.8 yrs (16-49), 89 individuals are males (59%), and 123 individuals are Caucasian of Non-Hispanic or Latino descent (82%). The Mayo Clinic Institutional Review Board approved the study and all subjects provided written informed consent. Data obtained from the Parkinson’s Progression Markers Initiative (PPMI) included 58 unrelated EOPD patients with an average age at onset of 44 yrs (33-49), 36 males (62%), and 54 are Caucasian of Non-Hispanic or Latino descent (93%).

### Whole Genome sequencing

Whole genome sequencing (WGS) was performed on the Illumina Hi-Seq platform using genomic DNA extracted from whole blood. The data was processed using the Mayo Genome GPS v4.0 pipeline. Functional annotations of variants were performed using ANNOVAR. Genotype calls with GQ < 10 and/or depth (DP) < 10 were set to missing, and variants with ED > 4 were removed from all subsequent analyses. For all analyses, only variants that passed Variant Quality Score Recalibration (VQSR) and with a call rate > 95% were considered, unless otherwise specified. The average coverage per exon of the *ATG9A* gene was over 100X, with a minimum coverage of 51X. The joint variant call file (gVCF) was then imported into Golden Helix SNP and Variation Suite (SVS) for further variant calling annotations. Using SVS, variants located within *ATG9A* were extracted and included annotation tracks: NCBI RefSeq Genes 109 Interim v2, NCBI dbSNP 149, and BROAD gnomAD Genomes Variant Frequencies 2.1.1.

### Data and Statistical Analysis

Data analysis and visualization were performed by GraphPad Prism version 9.3.1. All quantitative results are expressed as mean ± SD or SEM from at least three independent experiments. Statistical comparisons were performed using parametric t-test, one-way or two-way ANOVA as indicated in each figure legend. Significance levels are as follows: * p < 0.05, ** p < 0.005, *** p < 0.0005.

## ACKNOWLEDGMENT

We thank all patients who participate in research studies. We thank Susanne Heynen-Genel and Pedro Aza-Blanc from the Sanford-Prebys Medical institute in La Jolla, CA, for setting up and executing the genome-wide miRNA screen and validation screen, and Jenny M. Bredenberg for technical assistance.

## ABBREVIATIONS

ATG9A: Autophagy related protein 9a
CADD: combined annotation dependent depletion
CNV: copy number variant
DEG: differentially expressed gene
CCCP: Carbonyl cyanide m-chlorophenyl hydrazone
DMSO: dimethyl sulfoxide
EOPD: early-onset Parkinson’s disease
GFP: green fluorescent protein
KEGG: Kyoto Encyclopedia of Genes and Genomes
miRNA: microRNA
mRNA: messenger RNA
PINK1: PTEN-induced kinase 1
pS65-Ub: phosphorylated ubiquitin at serine 65
qRT-PCR: quantitative real-time reverse transcription PCR
siRNA: small interfering RNA
UTR: untranslated region

## REFERENCES

1. Shulman JM, De Jager PL, Feany MB. Parkinson’s disease: genetics and pathogenesis. Annu Rev Pathol. 2011;6:193–222.

2. Winklhofer KF, Haass C. Mitochondrial dysfunction in Parkinson’s disease. Biochim Biophys Acta. 2010 Jan;1802(1):29–44.

3. Geisler S, Holmstrom KM, Skujat D, et al. PINK1/Parkin-mediated mitophagy is dependent on VDAC1 and p62/SQSTM1. Nat Cell Biol. 2010 Feb;12(2):119–31.

4. Narendra DP, Jin SM, Tanaka A, et al. PINK1 is selectively stabilized on impaired mitochondria to activate Parkin. PLoS Biol. 2010 Jan 26;8(1):e1000298.

5. Exner N, Lutz AK, Haass C, et al. Mitochondrial dysfunction in Parkinson’s disease: molecular mechanisms and pathophysiological consequences. EMBO J. 2012 Jun 26;31(14):3038–62.

6. Kane LA, Lazarou M, Fogel AI, et al. PINK1 phosphorylates ubiquitin to activate Parkin E3 ubiquitin ligase activity. J Cell Biol. 2014 Apr 28;205(2):143–53.

7. Kazlauskaite A, Kondapalli C, Gourlay R, et al. Parkin is activated by PINK1-dependent phosphorylation of ubiquitin at Ser65. Biochem J. 2014 May 15;460(1):127–39.

8. Koyano F, Okatsu K, Kosako H, et al. Ubiquitin is phosphorylated by PINK1 to activate parkin. Nature. 2014 Jun 5;510(7503):162-6.

9. Kondapalli C, Kazlauskaite A, Zhang N, et al. PINK1 is activated by mitochondrial membrane potential depolarization and stimulates Parkin E3 ligase activity by phosphorylating Serine 65. Open Biol. 2012 May;2(5):120080.

10. Shiba-Fukushima K, Imai Y, Yoshida S, et al. PINK1-mediated phosphorylation of the Parkin ubiquitin-like domain primes mitochondrial translocation of Parkin and regulates mitophagy. Sci Rep. 2012;2:1002.

11. Iguchi M, Kujuro Y, Okatsu K, et al. Parkin-catalyzed ubiquitin-ester transfer is triggered by PINK1-dependent phosphorylation. J Biol Chem. 2013 Jul 26;288(30):22019–32.

12. Truban D, Hou X, Caulfield TR, et al. PINK1, Parkin, and Mitochondrial Quality Control: What can we Learn about Parkinson’s Disease Pathobiology? J Parkinsons Dis. 2017;7(1):13–29.

13. Lazarou M, Sliter DA, Kane LA, et al. The ubiquitin kinase PINK1 recruits autophagy receptors to induce mitophagy. Nature. 2015 Aug 20;524(7565):309-314.

14. Fiesel FC, Ando M, Hudec R, et al. (Patho-)physiological relevance of PINK1-dependent ubiquitin phosphorylation. EMBO Rep. 2015 Sep;16(9):1114–30.

15. Hou X, Fiesel FC, Truban D, et al. Age- and disease-dependent increase of the mitophagy marker phospho-ubiquitin in normal aging and Lewy body disease. Autophagy. 2018;14(8):1404–1418.

16. Ivatt RM, Sanchez-Martinez A, Godena VK, et al. Genome-wide RNAi screen identifies the Parkinson disease GWAS risk locus SREBF1 as a regulator of mitophagy. Proc Natl Acad Sci U S A. 2014 Jun 10;111(23):8494–9.

17. Hasson SA, Kane LA, Yamano K, et al. High-content genome-wide RNAi screens identify regulators of parkin upstream of mitophagy. Nature. 2013 Dec 12;504(7479):291-5.

18. McCoy MK, Kaganovich A, Rudenko IN, et al. Hexokinase activity is required for recruitment of parkin to depolarized mitochondria. Hum Mol Genet. 2014 Jan 1;23(1):145–56.

19. Hoshino A, Wang WJ, Wada S, et al. The ADP/ATP translocase drives mitophagy independent of nucleotide exchange. Nature. 2019 Nov;575(7782):375-379.

20. Kim J, Fiesel FC, Belmonte KC, et al. miR-27a and miR-27b regulate autophagic clearance of damaged mitochondria by targeting PTEN-induced putative kinase 1 (PINK1). Mol Neurodegener. 2016 Jul 26;11(1):55.

21. Fiesel FC, Moussaud-Lamodiere EL, Ando M, et al. A specific subset of E2 ubiquitin-conjugating enzymes regulate Parkin activation and mitophagy differently. J Cell Sci. 2014 Aug 15;127(Pt 16):3488–504.

22. Egan DF, Chun MG, Vamos M, et al. Small Molecule Inhibition of the Autophagy Kinase ULK1 and Identification of ULK1 Substrates. Mol Cell. 2015 Jul 16;59(2):285–97.

23. Fiesel FC, Caulfield TR, Moussaud-Lamodiere EL, et al. Structural and Functional Impact of Parkinson Disease-Associated Mutations in the E3 Ubiquitin Ligase Parkin. Hum Mutat. 2015 Aug;36(8):774–86.

24. Fiesel FC, James ED, Hudec R, et al. Mitochondrial targeted HSP90 inhibitor Gamitrinib-TPP (G-TPP) induces PINK1/Parkin-dependent mitophagy. Oncotarget. 2017 Dec 5;8(63):106233–106248.

25. Fiesel FC, Fricova D, Hayes CS, et al. Substitution of PINK1 Gly411 modulates substrate receptivity and turnover. Autophagy. 2022 Dec 5:1–22.

26. Yamano K, Kikuchi R, Kojima W, et al. Critical role of mitochondrial ubiquitination and the OPTN-ATG9A axis in mitophagy. J Cell Biol. 2020 Sep 7;219(9).

27. O’Loughlin T, Kruppa AJ, Ribeiro ALR, et al. OPTN recruitment to a Golgi-proximal compartment regulates immune signalling and cytokine secretion. J Cell Sci. 2020 Jun 15;133(12).

28. Katayama H, Kogure T, Mizushima N, et al. A sensitive and quantitative technique for detecting autophagic events based on lysosomal delivery. Chem Biol. 2011 Aug 26;18(8):1042–52.

29. Mailler E, Guardia CM, Bai X, et al. The autophagy protein ATG9A enables lipid mobilization from lipid droplets. Nat Commun. 2021 Nov 19;12(1):6750.

30. Mattera R, Park SY, De Pace R, et al. AP-4 mediates export of ATG9A from the trans-Golgi network to promote autophagosome formation. Proc Natl Acad Sci U S A. 2017 Dec 12;114(50):E10697–E10706.

31. Tai Y, Pu M, Yuan L, et al. miR-34a-5p regulates PINK1-mediated mitophagy via multiple modes. Life Sci. 2021 Jul 1;276:119415.

32. Qiu F, Wu Y, Xie G, et al. MiRNA-1976 Regulates the Apoptosis of Dopaminergic Neurons by Targeting the PINK1 Gene. J Integr Neurosci. 2023 Feb 22;22(2):45.

33. Yoo M, Choi DC, Murphy A, et al. MicroRNA-593-5p contributes to cell death following exposure to 1-methyl-4-phenylpyridinium (MPP(+)) by targeting PTEN-induced putative kinase 1 (PINK1). J Biol Chem. 2023 Apr 13:104709.

34. Tsujimoto T, Mori T, Houri K, et al. miR-155 inhibits mitophagy through suppression of BAG5, a partner protein of PINK1. Biochem Biophys Res Commun. 2020 Mar 12;523(3):707–712.

35. Liu G, Qian M, Chen M, et al. miR-21-5p Suppresses Mitophagy to Alleviate Hyperoxia-Induced Acute Lung Injury by Directly Targeting PGAM5. Biomed Res Int. 2020;2020:4807254.

36. Zuo W, Yan F, Liu Z, et al. miR-330 regulates Drp-1 mediated mitophagy by targeting PGAM5 in a rat model of permanent focal cerebral ischemia. Eur J Pharmacol. 2020 Aug 5;880:173143.

37. Hammond SM. An overview of microRNAs. Adv Drug Deliv Rev. 2015 Jun 29;87:3–14.

38. Tatura R, Kraus T, Giese A, et al. Parkinson’s disease: SNCA-, PARK2-, and LRRK2-targeting microRNAs elevated in cingulate gyrus. Parkinsonism Relat Disord. 2016 Dec;33:115–121.

39. Briggs CE, Wang Y, Kong B, et al. Midbrain dopamine neurons in Parkinson’s disease exhibit a dysregulated miRNA and target-gene network. Brain Res. 2015 Aug 27;1618:111–21.

40. Hoss AG, Labadorf A, Beach TG, et al. microRNA Profiles in Parkinson’s Disease Prefrontal Cortex. Front Aging Neurosci. 2016;8:36.

41. Goh SY, Chao YX, Dheen ST, et al. Role of MicroRNAs in Parkinson’s Disease. Int J Mol Sci. 2019 Nov 12;20(22).

42. Zhang Z, Zou J, Wang GK, et al. Uracils at nucleotide position 9-11 are required for the rapid turnover of miR-29 family. Nucleic Acids Res. 2011 May;39(10):4387–95.

43. Hwang HW, Wentzel EA, Mendell JT. A hexanucleotide element directs microRNA nuclear import. Science. 2007 Jan 5;315(5808):97-100.

44. Lian WS, Ko JY, Chen YS, et al. MicroRNA-29a represses osteoclast formation and protects against osteoporosis by regulating PCAF-mediated RANKL and CXCL12. Cell Death Dis. 2019 Sep 23;10(10):705.

45. Xu L, Xu Y, Jing Z, et al. Altered expression pattern of miR-29a, miR-29b and the target genes in myeloid leukemia. Exp Hematol Oncol. 2014;3:17.

46. Heid J, Cencioni C, Ripa R, et al. Age-dependent increase of oxidative stress regulates microRNA-29 family preserving cardiac health. Sci Rep. 2017 Dec 4;7(1):16839.

47. Matoba K, Kotani T, Tsutsumi A, et al. Atg9 is a lipid scramblase that mediates autophagosomal membrane expansion. Nat Struct Mol Biol. 2020 Dec;27(12):1185–1193.

48. Claude-Taupin A, Jia J, Bhujabal Z, et al. ATG9A protects the plasma membrane from programmed and incidental permeabilization. Nat Cell Biol. 2021 Aug;23(8):846–858.

49. Yamaguchi J, Suzuki C, Nanao T, et al. Atg9a deficiency causes axon-specific lesions including neuronal circuit dysgenesis. Autophagy. 2018;14(5):764–777.

50. Saitoh T, Fujita N, Hayashi T, et al. Atg9a controls dsDNA-driven dynamic translocation of STING and the innate immune response. Proc Natl Acad Sci U S A. 2009 Dec 8;106(49):20842–6.

51. Runwal G, Stamatakou E, Siddiqi FH, et al. LC3-positive structures are prominent in autophagy-deficient cells. Sci Rep. 2019 Jul 12;9(1):10147.

52. Maeda S, Yamamoto H, Kinch LN, et al. Structure, lipid scrambling activity and role in autophagosome formation of ATG9A. Nat Struct Mol Biol. 2020 Dec;27(12):1194–1201.

53. Zhou C, Ma K, Gao R, et al. Regulation of mATG9 trafficking by Src- and ULK1-mediated phosphorylation in basal and starvation-induced autophagy. Cell Res. 2017 Feb;27(2):184–201.

54. Itakura E, Kishi-Itakura C, Koyama-Honda I, et al. Structures containing Atg9A and the ULK1 complex independently target depolarized mitochondria at initial stages of Parkin-mediated mitophagy. J Cell Sci. 2012 Mar 15;125(Pt 6):1488–99.

55. Heo JM, Ordureau A, Swarup S, et al. RAB7A phosphorylation by TBK1 promotes mitophagy via the PINK-PARKIN pathway. Sci Adv. 2018 Nov;4(11):eaav0443.

56. Tan EHN, Tang BL. Rab7a and Mitophagosome Formation. Cells. 2019 Mar 8;8(3).

57. Consortium EP. The ENCODE (ENCyclopedia Of DNA Elements) Project. Science. 2004 Oct 22;306(5696):636-40.

58. Guardia CM, Tan XF, Lian T, et al. Structure of Human ATG9A, the Only Transmembrane Protein of the Core Autophagy Machinery. Cell Rep. 2020 Jun 30;31(13):107837.

59. Flinkman D, Hong Y, Gnjatovic J, et al. Regulators of proteostasis are translationally repressed in fibroblasts from patients with sporadic and LRRK2-G2019S Parkinson’s disease. NPJ Parkinsons Dis. 2023 Feb 6;9(1):20.

60. Pang HB, Braun GB, Friman T, et al. An endocytosis pathway initiated through neuropilin-1 and regulated by nutrient availability. Nat Commun. 2014 Oct 3;5:4904.

61. Kim J, Lee JE, Heynen-Genel S, et al. Functional genomic screen for modulators of ciliogenesis and cilium length. Nature. 2010 Apr 15;464(7291):1048-51.

62. Livak KJ, Schmittgen TD. Analysis of relative gene expression data using real-time quantitative PCR and the 2(-Delta Delta C(T)) Method. Methods. 2001 Dec;25(4):402–8.

63. Kalari KR, Nair AA, Bhavsar JD, et al. MAP-RSeq: Mayo Analysis Pipeline for RNA sequencing. BMC Bioinformatics. 2014 Jun 27;15:224.

64. Dobin A, Davis CA, Schlesinger F, et al. STAR: ultrafast universal RNA-seq aligner. Bioinformatics. 2013 Jan 1;29(1):15–21.

65. Liao Y, Smyth GK, Shi W. The Subread aligner: fast, accurate and scalable read mapping by seed-and-vote. Nucleic Acids Res. 2013 May 1;41(10):e108.

66. Wang L, Wang S, Li W. RSeQC: quality control of RNA-seq experiments. Bioinformatics. 2012 Aug 15;28(16):2184–5.

67. Robinson MD, McCarthy DJ, Smyth GK. edgeR: a Bioconductor package for differential expression analysis of digital gene expression data. Bioinformatics. 2010 Jan 1;26(1):139–40.

68. Watzlawik JO, Hou X, Fricova D, et al. Sensitive ELISA-based detection method for the mitophagy marker p-S65-Ub in human cells, autopsy brain, and blood samples. Autophagy. 2021 Sep;17(9):2613–2628.

69. McGeary SE, Lin KS, Shi CY, et al. The biochemical basis of microRNA targeting efficacy. Science. 2019 Dec 20;366(6472).

70. Karczewski KJ, Francioli LC, Tiao G, et al. The mutational constraint spectrum quantified from variation in 141,456 humans. Nature. 2020 May;581(7809):434-443.

71. Jumper J, Evans R, Pritzel A, et al. Highly accurate protein structure prediction with AlphaFold. Nature. 2021 Aug;596(7873):583-589.

