## Supplemental Information for "miRNA family miR-29 inhibits PINK1-PRKN dependent mitophagy via ATG9A"

[1] Department of Neuroscience, Mayo Clinic, Jacksonville, FL 32224, USA
[2] Revvity, Waltham, MA 02451, USA
[3] Division of Clinical Trials and Biostatistics, Mayo Clinic, Jacksonville, FL 32224, USA
[4] Department of Quantitative Health Science, Mayo Clinic, Jacksonville, FL 32224, USA
[5] Department of Quantitative Health Science, Mayo Clinic, Rochester, MN 55905, USA
[6] Department of Neurology, Mayo Clinic, Jacksonville, FL 32224, USA
[7] Neuroscience PhD Program, Mayo Clinic Graduate School of Biomedical Sciences, Jacksonville, FL 32224, USA
[#] present address: Department of Medical and Molecular Genetics, Stark Neurosciences Research Institute, Indiana University School of Medicine, Indianapolis, IN 46202, USA

#### FIGURE LEGENDS & TABLES

**Figure S1. High content imaging assay development.**

(**A**) GFP-PRKN cells were transfected with a non-targeting siRNA, a negative control miRNA, or transfection reagent only and assessed for the percentage of GFP-PRKN translocation positive cells at increasing CCCP concentrations. Shown is the mean ± SEM of duplicate wells from one independent experiment. (**B**) Representative images show classification of cell populations into GFP-PRKN translocation positive (red) versus negative cells (green). Border elements were excluded from the analysis.

**Figure S2. Validation of the RNAseq results.**

(**A**) Four genes with varying fold changes were chosen for general validation of RNA sequencing. (**B**) Cells were transfected with miR-29a and analyzed for fold change of mRNA for the chosen genes of COL4A1, HMGCR, CDC42, and VEGFA. Shown is the average fold change for three sets of experiments normalized to the control miRNA + SEM.

**Figure S3. Effects of miR-29 on pS65-Ub and PINK1.**

(**A**) HeLa cells transfected with miR-29a, miR-29b or miR-29c were analyzed using western blot for pS65-Ub signal after 2 h CCCP treatment. Shown is the mean normalized to GAPDH + SEM from at least four independent experiments. Statistical analysis was performed using two-way ANOVA with Dunnett's multiple comparisons test. (**B**) HeLa cells transfected with miR-29a, miR-29b, and miR-29c and analyzed for fold change of mRNA levels for PINK1 via qRT-PCR. Shown is the average fold change normalized to the control + SEM from at least four independent experiments. Statistical analysis was performed using one-way ANOVA with Dunnett's multiple comparisons test. (**C**) PINK1 levels were also analyzed using immunoblot analysis. Shown is the mean normalized to GAPDH + SEM. Statistical analysis was performed using two-way ANOVA with Dunnett's multiple comparisons test.

**Figure S4. Silencing of ATG9A reduces mitophagy.**

(**A**) Live cell high content mitolysosome analysis of HeLa GFP-PRKN MitoKeima cells transfected with scrambled siRNA control, ATG9A siRNA, or PINK1 siRNA treated with CCCP before imaging. Total area of acidic mitochondria over total mitochondria per cell was analyzed every two hours for the first six hours and then every four hours for a total of 18 hours. Shown is the mean of 18 replicate wells from one experiment ± SEM. Statistical analysis was performed using two-way ANOVA with Dunnett's multiple comparisons test. (**B**) Shown are representative 20x zoomed images of mitolysosomes for each of the timepoints for scrambled siRNA control (scr), ATG9A siRNA, and PINK1 siRNA transfected wells. Acidic mitochondria are colored red and neutral mitochondria are colored green. Scale bars represent 20 µm.

**Figure S5. ATG9A and miR-29a both increase lipid droplet number and size.**

(**A**) HeLa cells were transfected with either scrambled control siRNA or ATG9A siRNA and stained with the nuclear stain Hoechst and the lipid droplet probe BODIPY 493/503. Shown are representative images after treatment with 0, 2, and 4 h CCCP. (**B**) Shown is the mean ± SEM from three independent experiments of lipid droplet puncta number per cell and the mean lipid droplet area per cell. (**C**) HeLa cells were transfected with negative control miRNA or miR-29a and stained with Hoechst (blue) and the lipid droplet probe BODIPY 493/503 (green). Shown are representative images after treatment with 0, 2, and 4 h CCCP. (**D**) Shown is the mean + SEM from three independent experiments of lipid droplet puncta number per cell and the mean lipid droplet area per cell. Statistical analyses were performed using an unpaired t-test. (**E**) HeLa cells were stained with ATG9A, mitochondrial marker HSP60, and nuclear marker Hoechst. Shown are representative images after 0, 2, and 6 h CCCP treatment. Scale bars represent 10 µm.

**Table S1.** Imaging properties considered to find the best descriptor for PRKN translocation.

**Table S2.** Image analysis routine for miRNA screen.

**Table S3.** Final linear classifier to define PRKN translocation positive cells.

**Table S4.** Conserved 3'-UTR miR-29 binding sites and alignment of ATG9A to miR-29a/b/c.

**Table S5.** ATG9A species alignment at the R631 and S828 positions.

#### Figure S1

**A**

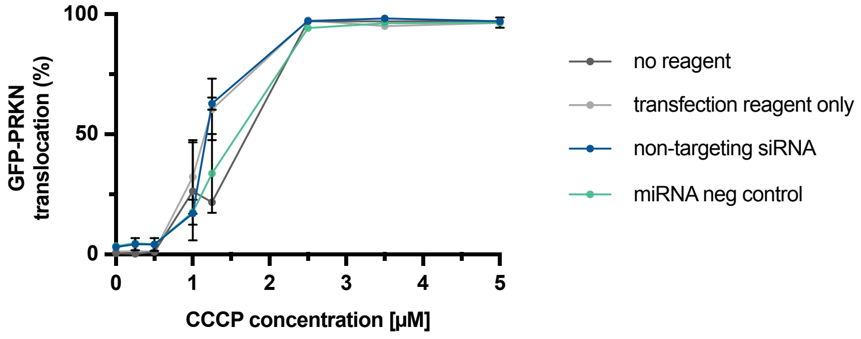

100
50
0
GFP-PRKN translocation (%)
0
1
2
3
4
5
CCCP concentration [µM]
no reagent
transfection reagent only
non-targeting siRNA
miRNA neg control


**B**

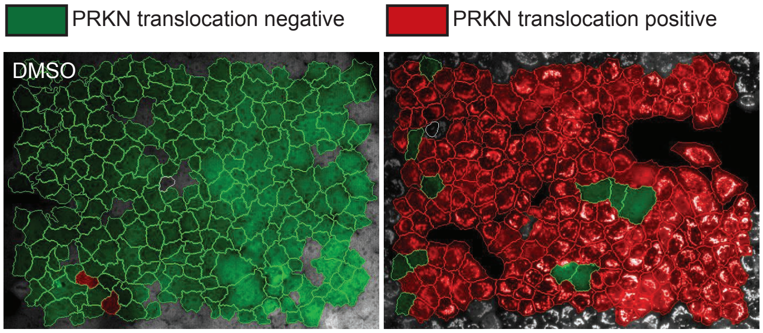

PRKN translocation negative
PRKN translocation positive
DMSO

#### Figure S2

A

| Gene | Fold Change |
|---|---|
| COL4A1 | 0.414 |
| HMGCR | 0.873 |
| CDC42 | 0.746 |
| VEGFA | 0.976 |

B

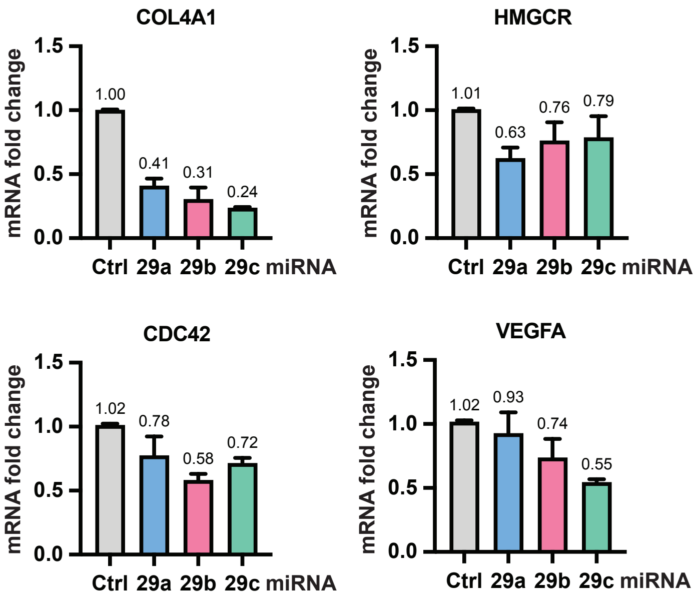

### Figure S3

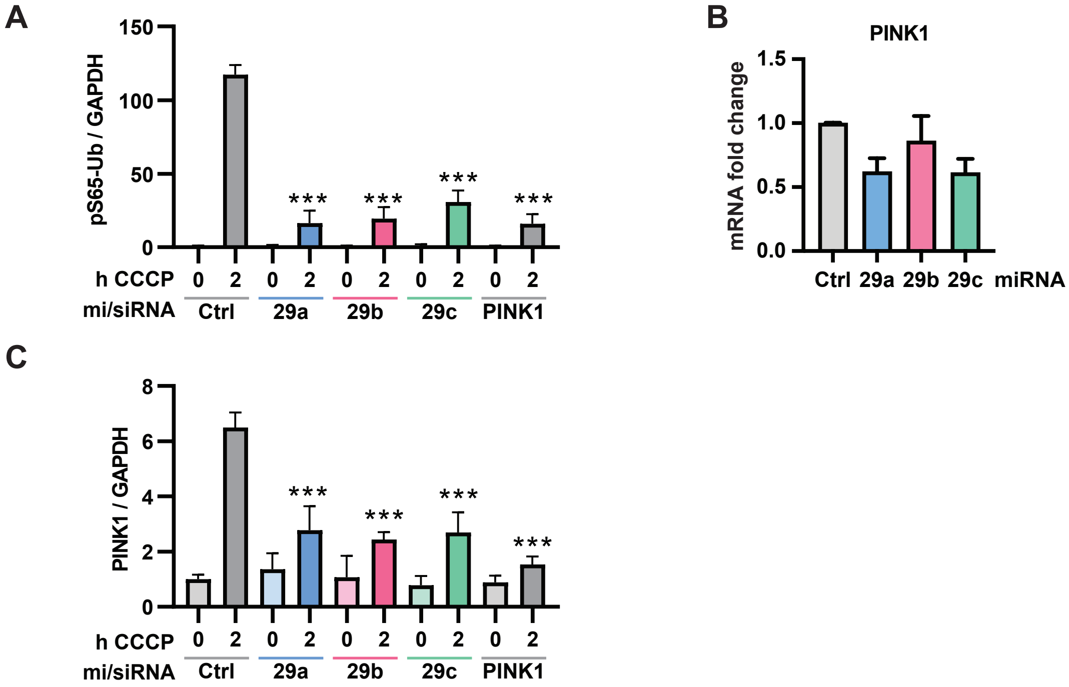
A
pS65-Ub / GAPDH
150
100
50
0
***
***
***
***
h CCCP 0 2 0 2 0 2 0 2 0 2
mi/siRNA Ctrl 29a 29b 29c PINK1
B
PINK1
mRNA fold change
1.5
1.0
0.5
0.0
Ctrl 29a 29b 29c miRNA
C
PINK1 / GAPDH
8
6
4
2
0
***
***
***
***
h CCCP 0 2 0 2 0 2 0 2 0 2
mi/siRNA Ctrl 29a 29b 29c PINK1

### Figure S4

A

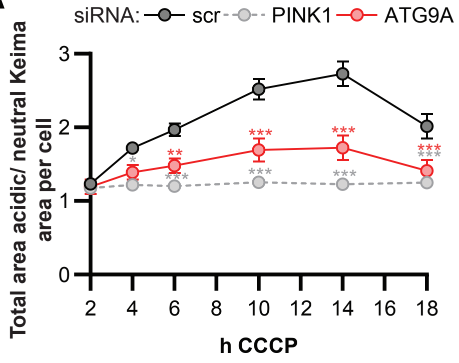
siRNA: scr PINK1 ATG9A
Total area acidic/ neutral Keima area per cell
3
2
1
0
2 4 6 8 10 12 14 16 18
h CCCP
*
**
***
***
***

B

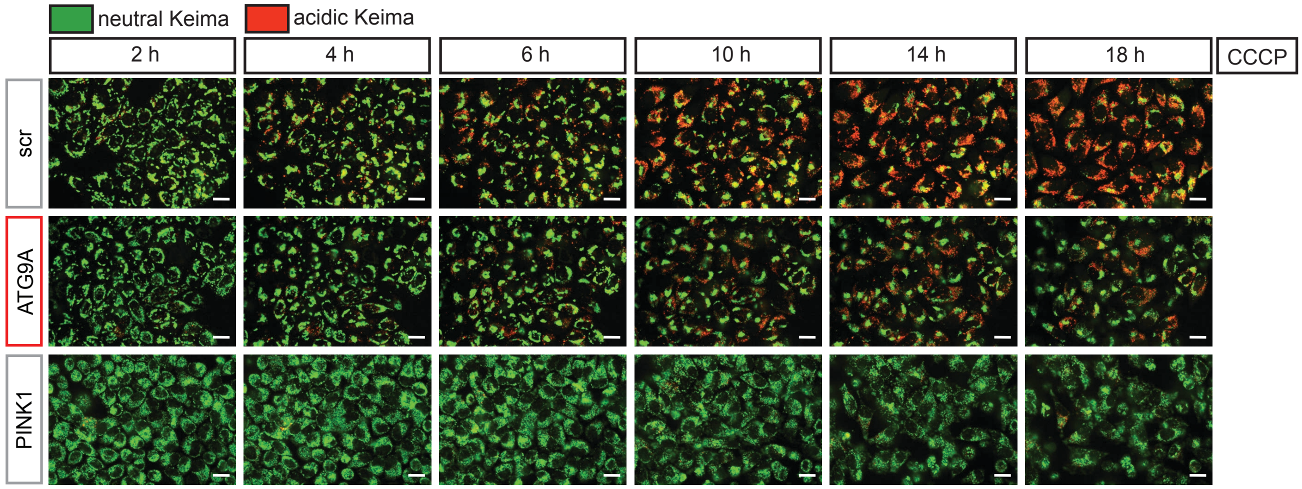
neutral Keima
acidic Keima
2 h
4 h
6 h
10 h
14 h
18 h
CCCP
scr
ATG9A
PINK1

#### Figure S5

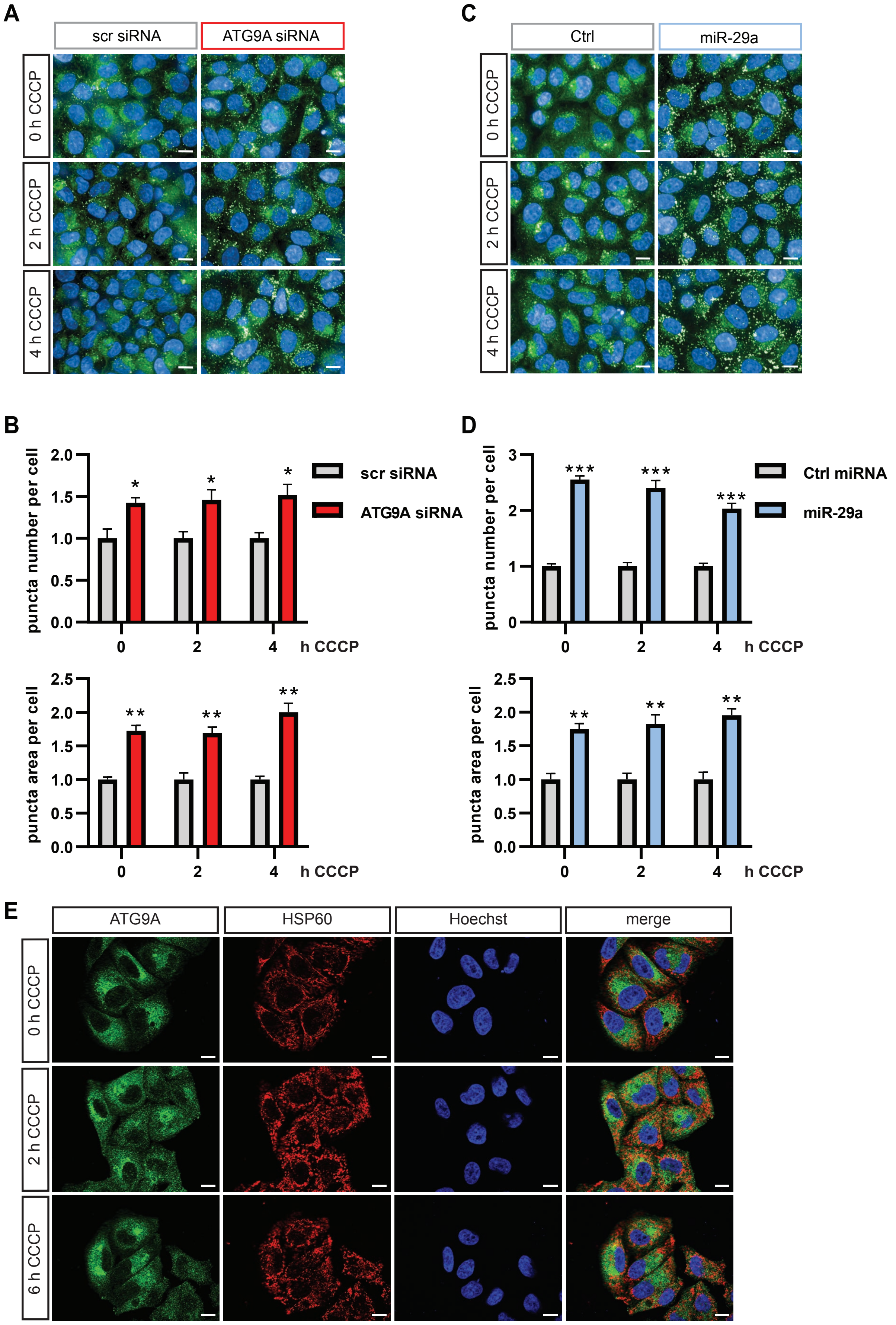

A
scr siRNA
ATG9A siRNA
0 h CCCP
2 h CCCP
4 h CCCP
C
Ctrl
miR-29a
0 h CCCP
2 h CCCP
4 h CCCP
B
puncta number per cell
2.0
1.5
1.0
0.5
0.0
0
2
4
h CCCP
scr siRNA
ATG9A siRNA
puncta area per cell
2.5
D
puncta number per cell
3
2
1
0
Ctrl miRNA
miR-29a
puncta area per cell
E
ATG9A
HSP60
Hoechst
merge
0 h CCCP
2 h CCCP
6 h CCCP

#### Table S1: Imaging properties considered to find the best descriptor for PRKN translocation

| 415 Properties Considered by PhenoLOGIC's Linear Classifier | |
|---|---|
| Cell Area [µm²] | a488Cell Radial Mean Ratio SER-Edge |
| Cell Roundness | a488Cell Profile 1/5 SER-Edge |
| Cell Ratio Width to Length | a488Cell Profile 2/5 SER-Edge |
| Nucleus Area [µm²] | a488Cell Profile 3/5 SER-Edge |
| Intensity Cell Exp2Cam4 Mean | a488Cell Profile 4/5 SER-Edge |
| Intensity Cell Calculated Image Mean | a488Cell Profile 5/5 SER-Edge |
| Intensity Cytoplasm Calculated Image Mean | calcCell Symmetry 02 |
| Intensity Cytoplasm Calculated Image StdDev | calcCell Symmetry 03 |
| Intensity Cytoplasm Calculated Image Maximum | calcCell Symmetry 04 |
| Intensity Nucleus Calculated Image Mean | calcCell Symmetry 05 |
| CalcImage Max Cyto/Mean Nuc | calcCell Symmetry 12 |
| Intensity Cell Exp1Cam1 Mean | calcCell Symmetry 13 |
| Intensity Cytoplasm Exp1Cam1 Mean | calcCell Symmetry 14 |
| Intensity Cytoplasm Exp1Cam1 StdDev | calcCell Symmetry 15 |
| Intensity Cytoplasm Exp1Cam1 Maximum | calcCell Threshold Compactness 30% |
| Intensity Nucleus Exp1Cam1 Mean | calcCell Threshold Compactness 40% |
| eGFP Max Cyto/Mean Nuc | calcCell Threshold Compactness 50% |
| a488Cell Symmetry 02 | calcCell Threshold Compactness 60% |
| a488Cell Symmetry 03 | calcCell Axial Small Length |
| a488Cell Symmetry 04 | calcCell Axial Length Ratio |
| a488Cell Symmetry 05 | calcCell Radial Mean |
| a488Cell Symmetry 12 | calcCell Radial Relative Deviation |
| a488Cell Symmetry 13 | calcCell Profile 1/5 |
| a488Cell Symmetry 14 | calcCell Profile 2/5 |
| a488Cell Symmetry 15 | calcCell Profile 3/5 |
| a488Cell Threshold Compactness 30% | calcCell Profile 4/5 |
| a488Cell Threshold Compactness 40% | calcCell Profile 5/5 |
| a488Cell Threshold Compactness 50% | calcCell Symmetry 02 SER-Bright |
| a488Cell Threshold Compactness 60% | calcCell Symmetry 03 SER-Bright |
| a488Cell Axial Small Length | calcCell Symmetry 04 SER-Bright |
| a488Cell Axial Length Ratio | calcCell Symmetry 05 SER-Bright |
| a488Cell Radial Mean | calcCell Symmetry 12 SER-Bright |
| a488Cell Radial Relative Deviation | calcCell Symmetry 13 SER-Bright |
| a488Cell Profile 1/5 | calcCell Symmetry 14 SER-Bright |
| a488Cell Profile 2/5 | calcCell Symmetry 15 SER-Bright |
| a488Cell Profile 3/5 | calcCell Threshold Compactness 30% SER-Bright |
| a488Cell Profile 4/5 | calcCell Threshold Compactness 40% SER-Bright |
| a488Cell Profile 5/5 | calcCell Threshold Compactness 50% SER-Bright |
| a488Cell Symmetry 02 SER-Bright | calcCell Threshold Compactness 60% SER-Bright |
| a488Cell Symmetry 03 SER-Bright | calcCell Axial Small Length SER-Bright |
| a488Cell Symmetry 04 SER-Bright | calcCell Axial Length Ratio SER-Bright |
| a488Cell Symmetry 05 SER-Bright | calcCell Radial Mean SER-Bright |
| a488Cell Symmetry 12 SER-Bright | calcCell Radial Relative Deviation SER-Bright |
| a488Cell Symmetry 13 SER-Bright | calcCell Radial Mean Ratio SER-Bright |
| a488Cell Symmetry 14 SER-Bright | calcCell Profile 1/5 SER-Bright |
| a488Cell Symmetry 15 SER-Bright | calcCell Profile 2/5 SER-Bright |
| a488Cell Threshold Compactness 30% SER-Bright | calcCell Profile 3/5 SER-Bright |
| a488Cell Threshold Compactness 40% SER-Bright | calcCell Profile 4/5 SER-Bright |
| a488Cell Threshold Compactness 50% SER-Bright | calcCell Profile 5/5 SER-Bright |
| a488Cell Threshold Compactness 60% SER-Bright | calcCell Symmetry 02 SER-Dark |
| a488Cell Axial Small Length SER-Bright | calcCell Symmetry 03 SER-Dark |
| a488Cell Axial Length Ratio SER-Bright | calcCell Symmetry 04 SER-Dark |
| a488Cell Radial Mean SER-Bright | calcCell Symmetry 05 SER-Dark |
| a488Cell Radial Relative Deviation SER-Bright | calcCell Symmetry 12 SER-Dark |
| a488Cell Radial Mean Ratio SER-Bright | calcCell Symmetry 13 SER-Dark |
| a488Cell Profile 1/5 SER-Bright | calcCell Symmetry 14 SER-Dark |
| a488Cell Profile 2/5 SER-Bright | calcCell Symmetry 15 SER-Dark |
| a488Cell Profile 3/5 SER-Bright | calcCell Threshold Compactness 30% SER-Dark |
| a488Cell Profile 4/5 SER-Bright | calcCell Threshold Compactness 40% SER-Dark |
| a488Cell Profile 5/5 SER-Bright | calcCell Threshold Compactness 50% SER-Dark |
| a488Cell Symmetry 02 SER-Dark | calcCell Threshold Compactness 60% SER-Dark |
| a488Cell Symmetry 03 SER-Dark | calcCell Axial Small Length SER-Dark |
| a488Cell Symmetry 04 SER-Dark | calcCell Axial Length Ratio SER-Dark |
| a488Cell Symmetry 05 SER-Dark | calcCell Radial Mean SER-Dark |
| a488Cell Symmetry 12 SER-Dark | calcCell Radial Relative Deviation SER-Dark |
| a488Cell Symmetry 13 SER-Dark | calcCell Radial Mean Ratio SER-Dark |
| a488Cell Symmetry 14 SER-Dark | calcCell Profile 1/5 SER-Dark |
| a488Cell Symmetry 15 SER-Dark | calcCell Profile 2/5 SER-Dark |
| a488Cell Threshold Compactness 30% SER-Dark | calcCell Profile 3/5 SER-Dark |
| a488Cell Threshold Compactness 40% SER-Dark | calcCell Profile 4/5 SER-Dark |
| a488Cell Threshold Compactness 50% SER-Dark | calcCell Profile 5/5 SER-Dark |

| | |
|---|---|
| a488Cell Threshold Compactness 60% SER-Dark | calcCell Symmetry 02 SER-Ridge |
| a488Cell Axial Small Length SER-Dark | calcCell Symmetry 03 SER-Ridge |
| a488Cell Axial Length Ratio SER-Dark | calcCell Symmetry 04 SER-Ridge |
| a488Cell Radial Mean SER-Dark | calcCell Symmetry 05 SER-Ridge |
| a488Cell Radial Relative Deviation SER-Dark | calcCell Symmetry 12 SER-Ridge |
| a488Cell Radial Mean Ratio SER-Dark | calcCell Symmetry 13 SER-Ridge |
| a488Cell Profile 1/5 SER-Dark | calcCell Symmetry 14 SER-Ridge |
| a488Cell Profile 2/5 SER-Dark | calcCell Symmetry 15 SER-Ridge |
| a488Cell Profile 3/5 SER-Dark | calcCell Threshold Compactness 30% SER-Ridge |
| a488Cell Profile 4/5 SER-Dark | calcCell Threshold Compactness 40% SER-Ridge |
| a488Cell Profile 5/5 SER-Dark | calcCell Threshold Compactness 50% SER-Ridge |
| a488Cell Symmetry 02 SER-Ridge | calcCell Threshold Compactness 60% SER-Ridge |
| a488Cell Symmetry 03 SER-Ridge | calcCell Axial Small Length SER-Ridge |
| a488Cell Symmetry 04 SER-Ridge | calcCell Axial Length Ratio SER-Ridge |
| a488Cell Symmetry 05 SER-Ridge | calcCell Radial Mean SER-Ridge |
| a488Cell Symmetry 12 SER-Ridge | calcCell Radial Relative Deviation SER-Ridge |
| a488Cell Symmetry 13 SER-Ridge | calcCell Radial Mean Ratio SER-Ridge |
| a488Cell Symmetry 14 SER-Ridge | calcCell Profile 1/5 SER-Ridge |
| a488Cell Symmetry 15 SER-Ridge | calcCell Profile 2/5 SER-Ridge |
| a488Cell Threshold Compactness 30% SER-Ridge | calcCell Profile 3/5 SER-Ridge |
| a488Cell Threshold Compactness 40% SER-Ridge | calcCell Profile 4/5 SER-Ridge |
| a488Cell Threshold Compactness 50% SER-Ridge | calcCell Profile 5/5 SER-Ridge |
| a488Cell Threshold Compactness 60% SER-Ridge | calcCell Symmetry 02 SER-Valley |
| a488Cell Axial Small Length SER-Ridge | calcCell Symmetry 03 SER-Valley |
| a488Cell Axial Length Ratio SER-Ridge | calcCell Symmetry 04 SER-Valley |
| a488Cell Radial Mean SER-Ridge | calcCell Symmetry 05 SER-Valley |
| a488Cell Radial Relative Deviation SER-Ridge | calcCell Symmetry 12 SER-Valley |
| a488Cell Radial Mean Ratio SER-Ridge | calcCell Symmetry 13 SER-Valley |
| a488Cell Profile 1/5 SER-Ridge | calcCell Symmetry 14 SER-Valley |
| a488Cell Profile 2/5 SER-Ridge | calcCell Symmetry 15 SER-Valley |
| a488Cell Profile 3/5 SER-Ridge | calcCell Threshold Compactness 30% SER-Valley |
| a488Cell Profile 4/5 SER-Ridge | calcCell Threshold Compactness 40% SER-Valley |
| a488Cell Profile 5/5 SER-Ridge | calcCell Threshold Compactness 50% SER-Valley |
| a488Cell Symmetry 02 SER-Valley | calcCell Threshold Compactness 60% SER-Valley |
| a488Cell Symmetry 03 SER-Valley | calcCell Axial Small Length SER-Valley |
| a488Cell Symmetry 04 SER-Valley | calcCell Axial Length Ratio SER-Valley |
| a488Cell Symmetry 05 SER-Valley | calcCell Radial Mean SER-Valley |
| a488Cell Symmetry 12 SER-Valley | calcCell Radial Relative Deviation SER-Valley |
| a488Cell Symmetry 13 SER-Valley | calcCell Radial Mean Ratio SER-Valley |
| a488Cell Symmetry 14 SER-Valley | calcCell Profile 1/5 SER-Valley |
| a488Cell Symmetry 15 SER-Valley | calcCell Profile 2/5 SER-Valley |
| a488Cell Threshold Compactness 30% SER-Valley | calcCell Profile 3/5 SER-Valley |
| a488Cell Threshold Compactness 40% SER-Valley | calcCell Profile 4/5 SER-Valley |
| a488Cell Threshold Compactness 50% SER-Valley | calcCell Profile 5/5 SER-Valley |
| a488Cell Threshold Compactness 60% SER-Valley | calcCell Symmetry 02 SER-Spot |
| a488Cell Axial Small Length SER-Valley | calcCell Symmetry 03 SER-Spot |
| a488Cell Axial Length Ratio SER-Valley | calcCell Symmetry 04 SER-Spot |
| a488Cell Radial Mean SER-Valley | calcCell Symmetry 05 SER-Spot |
| a488Cell Radial Relative Deviation SER-Valley | calcCell Symmetry 12 SER-Spot |
| a488Cell Radial Mean Ratio SER-Valley | calcCell Symmetry 13 SER-Spot |
| a488Cell Profile 1/5 SER-Valley | calcCell Symmetry 14 SER-Spot |
| a488Cell Profile 2/5 SER-Valley | calcCell Symmetry 15 SER-Spot |
| a488Cell Profile 3/5 SER-Valley | calcCell Threshold Compactness 30% SER-Spot |
| a488Cell Profile 4/5 SER-Valley | calcCell Threshold Compactness 40% SER-Spot |
| a488Cell Profile 5/5 SER-Valley | calcCell Threshold Compactness 50% SER-Spot |
| a488Cell Symmetry 02 SER-Spot | calcCell Threshold Compactness 60% SER-Spot |
| a488Cell Symmetry 03 SER-Spot | calcCell Axial Small Length SER-Spot |
| a488Cell Symmetry 04 SER-Spot | calcCell Axial Length Ratio SER-Spot |
| a488Cell Symmetry 05 SER-Spot | calcCell Radial Mean SER-Spot |
| a488Cell Symmetry 12 SER-Spot | calcCell Radial Relative Deviation SER-Spot |
| a488Cell Symmetry 13 SER-Spot | calcCell Radial Mean Ratio SER-Spot |
| a488Cell Symmetry 14 SER-Spot | calcCell Profile 1/5 SER-Spot |
| a488Cell Symmetry 15 SER-Spot | calcCell Profile 2/5 SER-Spot |
| a488Cell Threshold Compactness 30% SER-Spot | calcCell Profile 3/5 SER-Spot |
| a488Cell Threshold Compactness 40% SER-Spot | calcCell Profile 4/5 SER-Spot |
| a488Cell Threshold Compactness 50% SER-Spot | calcCell Profile 5/5 SER-Spot |
| a488Cell Threshold Compactness 60% SER-Spot | calcCell Symmetry 02 SER-Hole |
| a488Cell Axial Small Length SER-Spot | calcCell Symmetry 03 SER-Hole |
| a488Cell Axial Length Ratio SER-Spot | calcCell Symmetry 04 SER-Hole |
| a488Cell Radial Mean SER-Spot | calcCell Symmetry 05 SER-Hole |
| a488Cell Radial Relative Deviation SER-Spot | calcCell Symmetry 12 SER-Hole |
| a488Cell Radial Mean Ratio SER-Spot | calcCell Symmetry 13 SER-Hole |
| a488Cell Profile 1/5 SER-Spot | calcCell Symmetry 14 SER-Hole |
| a488Cell Profile 2/5 SER-Spot | calcCell Symmetry 15 SER-Hole |

| | |
|---|---|
| a488Cell Profile 3/5 SER-Spot | calcCell Threshold Compactness 30% SER-Hole |
| a488Cell Profile 4/5 SER-Spot | calcCell Threshold Compactness 40% SER-Hole |
| a488Cell Profile 5/5 SER-Spot | calcCell Threshold Compactness 50% SER-Hole |
| a488Cell Symmetry 02 SER-Hole | calcCell Threshold Compactness 60% SER-Hole |
| a488Cell Symmetry 03 SER-Hole | calcCell Axial Small Length SER-Hole |
| a488Cell Symmetry 04 SER-Hole | calcCell Axial Length Ratio SER-Hole |
| a488Cell Symmetry 05 SER-Hole | calcCell Radial Mean SER-Hole |
| a488Cell Symmetry 12 SER-Hole | calcCell Radial Relative Deviation SER-Hole |
| a488Cell Symmetry 13 SER-Hole | calcCell Radial Mean Ratio SER-Hole |
| a488Cell Symmetry 14 SER-Hole | calcCell Profile 1/5 SER-Hole |
| a488Cell Symmetry 15 SER-Hole | calcCell Profile 2/5 SER-Hole |
| a488Cell Threshold Compactness 30% SER-Hole | calcCell Profile 3/5 SER-Hole |
| a488Cell Threshold Compactness 40% SER-Hole | calcCell Profile 4/5 SER-Hole |
| a488Cell Threshold Compactness 50% SER-Hole | calcCell Profile 5/5 SER-Hole |
| a488Cell Threshold Compactness 60% SER-Hole | calcCell Symmetry 02 SER-Saddle |
| a488Cell Axial Small Length SER-Hole | calcCell Symmetry 03 SER-Saddle |
| a488Cell Axial Length Ratio SER-Hole | calcCell Symmetry 04 SER-Saddle |
| a488Cell Radial Mean SER-Hole | calcCell Symmetry 05 SER-Saddle |
| a488Cell Radial Relative Deviation SER-Hole | calcCell Symmetry 12 SER-Saddle |
| a488Cell Radial Mean Ratio SER-Hole | calcCell Symmetry 13 SER-Saddle |
| a488Cell Profile 1/5 SER-Hole | calcCell Symmetry 14 SER-Saddle |
| a488Cell Profile 2/5 SER-Hole | calcCell Symmetry 15 SER-Saddle |
| a488Cell Profile 3/5 SER-Hole | calcCell Threshold Compactness 30% SER-Saddle |
| a488Cell Profile 4/5 SER-Hole | calcCell Threshold Compactness 40% SER-Saddle |
| a488Cell Profile 5/5 SER-Hole | calcCell Threshold Compactness 50% SER-Saddle |
| a488Cell Symmetry 02 SER-Saddle | calcCell Threshold Compactness 60% SER-Saddle |
| a488Cell Symmetry 03 SER-Saddle | calcCell Axial Small Length SER-Saddle |
| a488Cell Symmetry 04 SER-Saddle | calcCell Axial Length Ratio SER-Saddle |
| a488Cell Symmetry 05 SER-Saddle | calcCell Radial Mean SER-Saddle |
| a488Cell Symmetry 12 SER-Saddle | calcCell Radial Relative Deviation SER-Saddle |
| a488Cell Symmetry 13 SER-Saddle | calcCell Radial Mean Ratio SER-Saddle |
| a488Cell Symmetry 14 SER-Saddle | calcCell Profile 1/5 SER-Saddle |
| a488Cell Symmetry 15 SER-Saddle | calcCell Profile 2/5 SER-Saddle |
| a488Cell Threshold Compactness 30% SER-Saddle | calcCell Profile 3/5 SER-Saddle |
| a488Cell Threshold Compactness 40% SER-Saddle | calcCell Profile 4/5 SER-Saddle |
| a488Cell Threshold Compactness 50% SER-Saddle | calcCell Profile 5/5 SER-Saddle |
| a488Cell Threshold Compactness 60% SER-Saddle | calcCell Symmetry 02 SER-Edge |
| a488Cell Axial Small Length SER-Saddle | calcCell Symmetry 03 SER-Edge |
| a488Cell Axial Length Ratio SER-Saddle | calcCell Symmetry 04 SER-Edge |
| a488Cell Radial Mean SER-Saddle | calcCell Symmetry 05 SER-Edge |
| a488Cell Radial Relative Deviation SER-Saddle | calcCell Symmetry 12 SER-Edge |
| a488Cell Radial Mean Ratio SER-Saddle | calcCell Symmetry 13 SER-Edge |
| a488Cell Profile 1/5 SER-Saddle | calcCell Symmetry 14 SER-Edge |
| a488Cell Profile 2/5 SER-Saddle | calcCell Symmetry 15 SER-Edge |
| a488Cell Profile 3/5 SER-Saddle | calcCell Threshold Compactness 30% SER-Edge |
| a488Cell Profile 4/5 SER-Saddle | calcCell Threshold Compactness 40% SER-Edge |
| a488Cell Profile 5/5 SER-Saddle | calcCell Threshold Compactness 50% SER-Edge |
| a488Cell Symmetry 02 SER-Edge | calcCell Threshold Compactness 60% SER-Edge |
| a488Cell Symmetry 03 SER-Edge | calcCell Axial Small Length SER-Edge |
| a488Cell Symmetry 04 SER-Edge | calcCell Axial Length Ratio SER-Edge |
| a488Cell Symmetry 05 SER-Edge | calcCell Radial Mean SER-Edge |
| a488Cell Symmetry 12 SER-Edge | calcCell Radial Relative Deviation SER-Edge |
| a488Cell Symmetry 13 SER-Edge | calcCell Radial Mean Ratio SER-Edge |
| a488Cell Symmetry 14 SER-Edge | calcCell Profile 1/5 SER-Edge |
| a488Cell Symmetry 15 SER-Edge | calcCell Profile 2/5 SER-Edge |
| a488Cell Threshold Compactness 30% SER-Edge | calcCell Profile 3/5 SER-Edge |
| a488Cell Threshold Compactness 40% SER-Edge | calcCell Profile 4/5 SER-Edge |
| a488Cell Threshold Compactness 50% SER-Edge | calcCell Profile 5/5 SER-Edge |
| a488Cell Threshold Compactness 60% SER-Edge | eGFP Cytoplasm Threshold Comp Threshold Compactness 30% |
| a488Cell Axial Small Length SER-Edge | eGFP Cytoplasm Threshold Comp Threshold Compactness 40% |
| a488Cell Axial Length Ratio SER-Edge | eGFP Cytoplasm Threshold Comp Threshold Compactness 50% |
| a488Cell Radial Mean SER-Edge | eGFP Cytoplasm Threshold Comp Threshold Compactness 60% |
| a488Cell Radial Relative Deviation SER-Edge | |

#### Table S2: Image analysis routine for miRNA screen

| Building Block | Purpose |
|---|---|
| 1. Find Image Region | Identifies the Whole Image as a Population and Region. |
| 2. Calculate Morphology | Calculates the Area [µm2] of the Whole Image |
| 3. Find Image Region | Finds Regions covered by Green (Exp1Cam1) |
| 4. Calculate Morphology | Calculates the Area [µm2] of the GreenArea |
| 5. Calculate Image | Normalized Green Channel minus Normalized Blue Channel, yields Green signal |
| 6. Find Nuclei | Finds Nuclei. Initial Population: DAPI_Pos |
| 7. Find Cytoplasm | Add Cell Regions to DAPI_Pos |
| 8. Select Population | Remove Cells that touch the image border. Output Population: Nuclei_Border_Filtered |
| 9. Calculate Morphology | Calculates the Area [µm2], Roundness and Length-to-Width Ratio of the Nuclei_Border_Filtered, Cell |
| 10. Calc. Morphology Properties | Calculates the Area [µm2] of the Nuclei_Border_Filtered, Nucleus Region |
| 11. Select Population | Selects Nuclei_Border_Filtered Objects based on Cell Roundness, Cell Area [µm2], and Nucleus Area |
| 12. Calculate Intensity Properties | Calculates Blue Intensity (Exp2Cam4) in the Cell Region of Nuclei |
| 13. Calculate Intensity Properties | Calculates ‘Calculated Image’ Intensity in Cell Region of Nuclei |
| 14. Calculate Intensity Properties | Calculates ‘Calculated Image’ Intensity in Cytoplasm Region of Nuclei |
| 15. Calculate Intensity Properties | Calculates ‘Calculated Image’ Intensity in Nucleus Region of Nuclei |
| 16. Calculate Properties | (Intensity Cytoplasm Calculated Image Max.) / (Intensity Nucleus Calculated Image Mean) |
| 17. Calculate Intensity Properties | Calculates Green Intensity (Exp1Cam1) in the Cell Region of Nuclei |
| 18. Calculate Intensity Properties | Calculates Green Intensity (Exp1Cam1) in the Cytoplasm Region of Nuclei |
| 19. Calculate Intensity Properties | Calculates Green Intensity (Exp1Cam1) in the Nucleus Region of Nuclei |
| 20. Calculate Properties | (Intensity Cytoplasm Exp1Cam1 Max.) / (Intensity Nucleus Exp1Cam1 Mean) |
| 21. Calculate Morphology | STAR Morphology with Texture Features, Exp1Cam1 Channel, Cell Region of Nuclei |
| 22. Calculate Morphology | STAR Morphology with Texture Features, ‘Calculated Image’ Channel, Cell Region of Nuclei |
| 23. Calculate Morphology | Threshold Compactness, Exp1Cam1(Green) of Cytoplasm Region of Nuclei |
| 24. Select Population | Linear Classifier on Nuclei, 415 properties considered |
| | Output Population A : DMSO |
| | Output Population B : 3.5_CCCP |
| | Output Population C : InBetween |
| 25. Calculate Properties | By Related Populations |
| | Relates the Area [µm2] of the Population pGreenAreas to the Population Whole Image |
| 26. Calculate Properties | By Related Populations |
| | Relates the Cell Area [µm2] of the Population Nuclei to the Population Whole Image |
| 27. Calculate Properties | Percentage of the Whole Image Area [µm2] covered by pGreenAreas. |
| | ((rGreenArea Area-Sum per Object) / (Whole Image Region Area) )*100 |
| 28. Calculate Properties | Percentage of the Whole Image Area [µm2] covered by segmented Cells. |
| | ((Cell Area-Sum per Object) / (Whole Image Region Area) )*100 |
| 29. Select Population | Flags fields of view less than 50% confluent. |
| 30. Define Results | Specifies data to be included in the final output table. |

##### Table S3: Final linear classifier to define PRKN translocation positive cells

Goodness: 5.21

Offset: -7.42822

| Properties (ordered by relevance) | Linear Coefficient |
|---|---|
| a488Cell Threshold Compactness 40% | -22.3388 |
| a488Cell Profile 4/5 SER-Dark | -135.71 |
| a488Cell Profile 3/5 SER-Bright | 91.583 |
| eGFP Cytoplasm Threshold Comp Threshold Compactness 40% | 22.9283 |
| a488Cell Threshold Compactness 50% | -14.4039 |
| a488Cell Profile 3/5 SER-Edge | 32.0958 |
| calcCell Profile 3/5 | 2.23318 |
| calcCell Threshold Compactness 60% SER-Edge | 9.08955 |
| a488Cell Profile 3/5 SER-Saddle | 75.3239 |
| eGFP Max Cyto/Mean Nuc | 0.945125 |
| calcCell Radial Mean SER-Ridge | -0.167702 |
| a488Cell Radial Mean Ratio SER-Valley | -5.2088 |

**Table S4: conserved 3'-UTR miR-29 binding sites and alignment of ATG9A to miR-29a/b/c.**

| miRNA | 3'-UTR Binding Site | Predicted Alignment of ATG9A (top) to miR-29 (bottom) | Cumulative weighted context++ score | Aggregate $P_{CT}$ |
|---|---|---|---|---|
| **miR-29a** | 251-258 | 5' ...CCUUGGCUCAGAGUGUGGUGCUA...<br>\|\|\|\|\|\|\|<br>3' AUUGGCUAAAGUCUACCACGAU | -0.28 | 0.88 |
| **miR-29b** | 251-258 | 5' ...CCUUGGCUCAGAGUGUGGUGCUA...<br>\|\|\|\|\|\|\|<br>3' UUGUGACUAAAGUUUACCACGAU | -0.26 | 0.88 |
| **miR-29c** | 251-258 | 5' ...CCUUGGCUCAGAGUGUGGUGCUA...<br>\|\|\|\|\|\|\|<br>3' AUUGGCUAAAGUUUACCACGAU | -0.26 | 0.88 |

Data obtained from TargetScan version 7.1.

**Table S5: ATG9A species alignment at the R631 and S828 positions**

| Species | R631 | S828 |
|---|---|---|
| *Homo sapiens* | AGSSCRGPPLP | VPEEGSEDELP |
| *Pongo abelii* | AGSSCRGPPLP | VPEEGSEDELP |
| *Mus musculus* | AGSSCRGPSLS | VPEEGSEDELP |
| *Sus scrofa* | AGSSCRGPPLP | VPEEGSEDELP |
| *Rattus norvegicus* | AGSSCRGPPLS | VPEEGSEDELP |
| *Bos taurus* | AGSSCRGPPLP | VPEEGSEDELP |

Data obtained from UniProt. R631 and S828 are highlighted in red.
